# Selective autophagy promotes bacterial immunity under warming through NBR1-dependent regulation of ABI5

**DOI:** 10.64898/2026.07.01.735842

**Authors:** Lara Anzardi Ruffino, Joaquín Suárez, Anahí Mara Yáñez Santos, Virginia Laura Lobatto, Verónica Sofía Mary, Martín Gustavo Theumer, María Clara Mesquida Nardini, Nicolás Miguel Cecchini, Hernán Ramiro Lascano, Ignacio Lescano López I

## Abstract

Elevated temperatures compromise plant immunity and increase susceptibility to bacterial pathogens through extensive reprogramming of hormone signaling pathways. Although autophagy contributes to both stress adaptation and pathogen defense, its role in hormone-dependent immune regulation under warm conditions remains unclear. Here, we investigated the contribution of NBR1 (NEIGHBOR OF BRCA1 GENE 1)-mediated selective autophagy to Arabidopsis immunity against *Pseudomonas cannabina* pv. *alisalensis* at elevated temperature. Bacterial infection under warming enhanced autophagic flux and promoted NBR1 turnover, indicating increased autophagic activity. Analysis of *atg5* and *nbr1* mutants, and NBR1-overexpressing lines, demonstrated that both core autophagy and NBR1-mediated selective autophagy contribute to bacterial immunity under warm conditions. Hormone and gene expression analyses indicated that NBR1 negatively regulates abscisic acid (ABA)-associated transcriptional responses during infection, while salicylic acid signaling was largely unaffected. Mechanistically, NBR1 physically associated with the ABA-responsive transcription factor ABI5 (ABA INSENSITIVE 5) and promoted its autophagy-dependent turnover *in planta*. ABI5 turnover was strongly reduced under warm conditions, leading to its accumulation in *nbr1* and *atg5* plants. Consistent with a functional role for ABI5 in this phenotype, genetic disruption of ABI5 largely reversed the increased susceptibility of *nbr1* mutants at elevated temperature, whereas ABI5 overexpression increased susceptibility to bacterial infection. Together, our results identify NBR1-mediated selective autophagy as a regulatory mechanism that restrains ABA-associated susceptibility through the autophagy-dependent turnover of ABI5. These findings reveal a previously unrecognized connection between selective autophagy and ABA-dependent immune regulation and identify NBR1-mediated ABI5 turnover as a temperature-dependent mechanism that prevents stronger bacterial susceptibility under warm conditions.

## INTRODUCTION

Plant diseases represent a primary cause of global crop losses, and their impact is expected to intensify with climate change (Savary et al. 2019, Velásquez et al., 2018). Plants rely on layered immune responses, with plasma membrane-localized pattern-recognition receptors (PRRs) detecting conserved microbial patterns to activate pattern-triggered immunity (PTI), and intracellular immune receptors recognizing pathogen effectors, many of which are delivered into host cells through specialized secretion systems (e.g. bacterial type III secretion system, T3SS), to activate effector-triggered immunity (ETI). Environmental factors critically influence the outcome of plant–pathogen interactions, with disease outbreaks frequently arising under suboptimal growth conditions. Because biotic and abiotic stress responses often involve competing for physiological programs, plants must maintain a dynamic equilibrium between growth, stress acclimation, and immunity (Huot et al., 2014; Karasov et al., 2017).

High temperatures shift the balance between growth and defense in favor of growth, thereby compromising immune responses (Kim et al., 2022). In Arabidopsis, moderately elevated temperatures (28–30 °C) trigger thermomorphogenesis, characterized by growth-related responses that often occur at the expense of defense (Casal and Balasubramanian, 2019). Studies in Arabidopsis and other plant species have shown that elevated temperatures attenuate immune responses through multiple mechanisms, including suppression of salicylic acid (SA)-dependent defenses and transcriptional and hormonal reprogramming (Zhu et al., 2010; Alcázar and Parker, 2011; Huot et al., 2017; Velásquez et al., 2018). In Arabidopsis, thermosensory pathways involving phytochrome B (phyB) and PHYTOCHROME INTERACTING FACTOR 4 (PIF4) have emerged as important regulators of the growth–defense balance under elevated temperature conditions, linking thermomorphogenic growth responses with changes in immune signaling (Gangappa and Kumar, 2017; Casal and Balasubramanian, 2019). Consequently, plants exposed to elevated temperatures often display enhanced susceptibility to bacterial pathogens, highlighting the need to better understand the molecular mechanisms that coordinate temperature perception, growth responses, and immunity.

Warm temperature reprograms phytohormone signaling, promoting growth-related pathways such as auxin and gibberellin while reshaping defense-associated hormonal networks (Quint et al., 2016; Casal and Balasubramanian, 2019). In Arabidopsis, transcriptomic analyses of *Pseudomonas syringae* pv. *tomato* (*Pst*)-infected plants revealed that elevated temperature alters the expression of numerous hormone-responsive genes, including components of the salicylic acid (SA), jasmonic acid (JA), and abscisic acid (ABA) pathways (Huot et al., 2017). Notably, the ABA biosynthetic gene *NCED3* is more strongly induced during infection under warm conditions, accompanied by increased accumulation of ABA. These changes are particularly relevant because ABA regulates multiple defense-associated processes, including stomatal closure, responses to water stress, and crosstalk with immune signaling pathways (Cutler et al., 2010; Yoshida et al., 2019). Moreover, ABA signaling can antagonize SA-dependent defenses, thereby promoting susceptibility to biotrophic and hemibiotrophic pathogens (De Torres-Zabala et al., 2007; Cao et al., 2011; Mine et al., 2017). Together, these observations suggest that ABA may represent an important regulatory node linking temperature perception and immune responses during pathogen infection. However, the mechanisms that integrate temperature-dependent hormonal reprogramming with immune regulation remain poorly understood.

Autophagy has emerged as a central mechanism by which plants maintain cellular homeostasis and adapt to changing environmental conditions. During this process, cytoplasmic material is engulfed into double-membrane vesicles known as autophagosomes and delivered to the vacuole for degradation and recycling. Autophagosome biogenesis depends on a conserved set of AUTOPHAGY-RELATED (ATG) proteins that mediate cargo sequestration, vesicle formation, and vacuolar delivery (Marshall and Vierstra, 2026). While autophagy can mediate bulk degradation of cytoplasmic components, increasing evidence indicates that many substrates are targeted through selective autophagy pathways mediated by cargo receptors (Stephani and Dagdas, 2020). A major mediator of selective autophagy in plants is NEIGHBOR OF BRCA1 GENE 1 (NBR1), a multifunctional cargo receptor that contains ubiquitin-binding and ATG8-interacting domains, enabling the recognition of ubiquitinated substrates and their delivery to autophagosomes for subsequent vacuolar degradation (Svenning et al., 2011). NBR1-dependent selective autophagy participates in the removal of protein aggregates (aggrephagy), the turnover of regulatory proteins involved in stress adaptation and development and in the clearance of pathogen-derived components (xenophagy) (Svenning et al., 2011; Stephani and Dagdas, 2020; Marshall and Vierstra, 2026). Consistent with its role as a cargo receptor, NBR1 accumulates in autophagy-deficient mutants such as *atg5*, reflecting its dependence on a functional autophagy pathway for turnover (Üstün et al., 2018). In addition to its functions in proteostasis, NBR1 contributes to plant responses to abiotic stresses, including recovery from heat-induced proteotoxic stress, and has emerged as an important regulator of immunity against diverse pathogens (Zhou et al., 2013; Hafrén et al., 2017; Üstün et al., 2018). For example, during *Pseudomonas syringae* infection, NBR1 restricts the development of HopM1-dependent water-soaked lesions and limits disease progression, demonstrating that selective autophagy can counteract bacterial virulence strategies (Üstün et al., 2018). Similarly, NBR1/Joka2 targets the Xanthomonas effector XopL, thereby limiting bacterial virulence and revealing that autophagy can directly counteract effector-mediated susceptibility (Leong et al., 2022). In addition, core autophagy components have been implicated in both pro- and antibacterial functions during *P. syringae* infection, highlighting the context-dependent role of autophagy in plant immunity (Hofius et al., 2009; Lenz et al., 2011; Üstün et al., 2018). Together, these findings indicate that distinct autophagy pathways, including NBR1-mediated selective autophagy, contribute to plant defense against bacterial pathogens.

Beyond its role in cellular homeostasis, autophagy also modulates hormone signaling by directly targeting specific regulatory proteins. Several hormone-associated components have been identified as autophagy substrates or interactors, highlighting selective autophagy as an important mechanism for fine-tuning hormonal responses. For example, ARF7 undergoes NBR1-dependent selective autophagic turnover, contributing to auxin homeostasis and root development (Ebstrup et al., 2024). Moreover, Arabidopsis NBR1 physically interacts with ABI3, ABI4, and ABI5 (ABA Insensitive) *in planta* (Tarnowski et al., 2020), basic leucine zipper (bZIP) transcription factors that activate ABA-responsive genes involved in stress adaptation, development, and stomatal regulation (Skubacz et al., 2016; Chandrasekaran et al., 2020; Cheng et al., 2026). This link is particularly relevant during *Pseudomonas syringae* infection, where ABA promotes HopM1-dependent water-soaked lesion formation, while enhanced NBR1 activity attenuates these disease symptoms (De Torres-Zabala et al., 2007; Üstün et al., 2018). Together, these observations suggest that NBR1-mediated selective autophagy may modulate ABA-dependent responses by regulating key signaling components such as ABI5.

Here, we investigated the contribution of NBR1-dependent selective autophagy to plant immunity at elevated temperature. We show that bacterial infection under warm conditions enhances autophagic flux and promotes NBR1 turnover, and that both NBR1 and core autophagy contribute to restricting bacterial proliferation. We further demonstrate that NBR1 negatively regulates ABA signaling, physically associates with ABI5 *in planta*, and promotes its autophagy-dependent turnover during infection. Finally, genetic analyses reveal that ABI5 contributes to the enhanced susceptibility of *nbr1* plants under warming conditions. Together, our findings identify NBR1-mediated selective autophagy as a regulatory mechanism linking temperature-responsive immunity and ABA signaling through the control of ABI5 homeostasis.

## MATERIALS AND METHODS

### Plant material and growth conditions

*Arabidopsis thaliana* (L.) Heynh. accession Columbia-0 (Col-0) was used as the wild-type (WT) background in this study. The T-DNA insertion mutants *nbr1* (GABI_246H08; Zhou et al. (2013)), *atg5* (SAIL_129B07; Inoue et al. (2006)), and *abi5-8* (SALK_013163; Zou et al. (2013)), as well as the 35S:GFP-ATG8a reporter line (N39996; Thompson et al. (2005)), were obtained from the Arabidopsis Biological Resource Center (ABRC, Ohio State University, USA). Seeds of the mCherry–YFP–NBR1 reporter line (Svenning et al., 2011) were kindly provided by Germán Robert (UDEA-CONICET). Seeds of the *abi5-8* line (Zou et al., 2013) were kindly provided by Elina Welchen (Instituto de Agrobiotecnología del Litoral (CONICET-UNL), Argentina). Seeds were imbibed in water and stratified at 4 °C for 2–3 days before sowing in a soil: vermiculite mixture (1:2, v/v). Plants were grown under a 12 h light/12 h dark photoperiod at 22 °C and a light intensity of 120–130 μmol m⁻² s⁻¹ unless otherwise indicated.

### Bacterial infection assays and temperature treatments

Virulent *Pseudomonas cannabina* pv. *alisalensis* (*Pma*DG3; formerly *P. syringae* pv. *maculicola*; Baltrus et al. (2011), *P. syringae* pv. *tomato* DC3000 (*Pst*; Ritter and Dangl (1996), the type III secretion-deficient mutant *Pst-*Δ*hrcC* (Hauck et al., 2003), and *Pst* expressing AvrRpt2 (*Pst-AvrRpt2*) were grown on King’s B agar plates supplemented with appropriate antibiotics for 2–3 days at 28 °C. Bacterial suspensions were prepared in 10 mM MgCl₂ and adjusted to the desired concentration based on OD₆₀₀ measurements.

To evaluate the effect of elevated temperature on bacterial infection, leaves from 4–5-week-old WT or transgenic plants were inoculated by syringe infiltration (5 × 10 CFU ml⁻¹). Mock-treated plants were inoculated with 10 mM MgCl₂ alone. Following inoculation, plants were maintained at either 22 °C or 29 °C for 1 or 2 days post-inoculation (dpi), depending on the experiment. Fully expanded leaves (leaves 7–9) were used for all infection assays.

Bacterial growth was quantified at 2 dpi according to a modified protocol from Miranda de la Torre et al. (2023). Briefly, for each treatment, at least three infected leaves from six to eight independent plants were collected. Two leaf discs (12.56 mm² each) were excised from each leaf, homogenized in 10 mM MgCl₂, serially diluted, and plated on LB agar supplemented with appropriate antibiotics. Colonies were counted after 2–3 days of incubation at 28 °C, and bacterial growth was expressed as colony-forming units (CFU) per cm² of leaf tissue. To estimate temperature-dependent changes in susceptibility, a susceptibility index (SI) was calculated as the ratio between bacterial titers at 29 °C and the mean bacterial titers at 22 °C for the corresponding genotype.

### Gene expression analysis

Leaf tissue was homogenized in liquid air using a plastic pestle, and total RNA was extracted using 1 ml of BIO-ZOL reagent (PB-L, Argentina) according to the manufacturer’s instructions. Total RNA (500 ng) was reverse-transcribed using oligo(dT) primers and M-MLV reverse transcriptase (K1600, Inbio Highway) according to the manufacturer’s instructions.

Transcript levels were analyzed by reverse transcription quantitative PCR (RT-qPCR) as previously described (Lescano Lopez et al., 2024), with modifications. Reactions were performed in a final volume of 15 μl using 1:20 diluted cDNA, DreamTaq DNA polymerase (EP0702, Thermo Fisher Scientific), and EvaGreen Dye (Biotium) in a Bioer qPCR instrument. The amplification program consisted of 94 °C for 2 min, followed by 45 cycles of 94 °C for 30 s, 60 °C for 20 s, and 72 °C for 20 s, followed by a melting curve analysis from 60 °C to 95 °C at 0.2 °C s⁻¹. Primer sequences are listed in Supplemental Table 1. Raw fluorescence data obtained using Gene 9660 V1.0.13 software (Bioer, 2011) were baseline-corrected, and the initial target quantity (N₀) was estimated using LinRegPCR 2021.1 (Ruijter et al., 2009). Relative transcript abundance was calculated by normalizing N₀ values to those of the reference gene *PP2A-A3* (PROTEIN PHOSPHATASE 2A SUBUNIT A3; AT1G13320).

To estimate the effect of elevated temperature on gene expression, the ratio between transcript levels at 29 °C and the mean transcript levels at 22 °C was calculated for each genotype and treatment.

### Hormone quantification

Phytohormone quantification was performed by liquid chromatography coupled to tandem mass spectrometry (LC-MS/MS) as previously described (Pettinari et al., 2022). Briefly, 50–100 mg of frozen tissue was ground to a fine powder in liquid nitrogen and homogenized in 500 μl of 1-propanol/H₂O/HCl (2:1:0.002, v/v/v). Samples were incubated for 30 min at 4 °C with agitation, followed by the addition of 1 ml dichloromethane (CH₂Cl₂) and a second incubation for 30 min at 4 °C with agitation.

Samples were centrifuged at 13,000 × g for 5 min, and approximately 1 ml of the lower organic phase was recovered and evaporated to dryness. Dried extracts were resuspended in 200 μl 50:50 (v/v) methanol (HPLC grade)-water solution containing 0.1 % formic acid and filtered prior to analysis.

Phytohormone analyses were carried out using a Waters Xevo TQ-S Micro LC-MS/MS system (Waters, Milford, MA, USA) equipped with a quaternary pump (Acquity UPLC H-Class, Waters), an autosampler (Acquity UPLC H-Class, Waters), and a reverse-phase column (Waters BEH C18, 1.7 µm, 2.1 × 50 mm). The mobile phase consisted of water containing 0.1% formic acid (A) and methanol containing 0.1% CH_2_O_2_ (B), with a flow rate of 0.25 ml min⁻¹. The initial concentration of solvent B was maintained at 40% for 0.5 min and then linearly increased to 100% at 3 min.

For metabolite identification and quantification, the UPLC system was coupled to a Xevo TQ-S micro triple quadrupole mass spectrometer (Waters) operating with an electrospray ionization (ESI) source. Data acquisition and processing were performed using MassLynx software version 4.1. Mass spectra were recorded in both positive and negative ionization modes. The mass transition (m/z) values used for each metabolite were as follows: SA: 137.0 > 93.0 and 137.0 > 65.0; IAA: 176.0 > 130.0 and 176.0 > 103.0; and ABA, 263.0 > 153.0 and 263.0 > 219.0. Phytohormone concentrations were calculated using linear calibration curves and expressed as ng of phytohormone per g of fresh tissue weight.

### Generation of transgenic lines expressing ABI5 fusion proteins

To generate transgenic lines expressing ABI5 fusion proteins, WT, *nbr1*, and *atg5* plants were transformed with pSuper:ABI5–3×Flag or pSuper:ABI5–GFP constructs (Qi et al., 2020), kindly provided by Professor Jigang Li (China Agricultural University), using *Agrobacterium tumefaciens* strain GV3101 and a modified floral-dip protocol based on Clough and Bent (1998). Transgenic plants from T1 to T3 generations were selected on 0.5× MS agar plates supplemented with hygromycin B (15 μg ml⁻¹), and homozygous lines were identified for subsequent analyses. Expression of ABI5–Flag and ABI5–GFP fusion proteins was verified by immunoblot analysis using anti-Flag or anti-GFP antibodies, respectively, as described below. Representative homozygous T3 lines were selected for subsequent analyses.

### Immunoblot analysis

Six leaf discs (12.56 mm² each) were collected and ground in liquid nitrogen using plastic pestles. Proteins were extracted directly in 100 μl of 1× SDS-PAGE loading buffer as previously described (Svenning et al., 2011). Samples were boiled at 95 °C for 5 min, centrifuged at 12,000 rpm, and 20 μl of supernatant was loaded per lane onto SDS-PAGE gels. Protein separation, transfer, and immunodetection were performed essentially as described by Robert et al. (2025), with minor modifications.

Following electrophoresis, proteins were transferred onto PVDF membranes, and equal loading was verified by Ponceau S staining. Membranes were blocked for 1 h in TBS-T buffer (50 mM Tris-HCl pH 7.5, 150 mM NaCl, 0.1% Tween-20) supplemented with 5% skim milk and incubated overnight at 4 °C with primary antibodies diluted 1:1000. The following primary antibodies were used: rabbit anti-GFP (AS20, Agrisera), rabbit anti-mCherry (AS15 4085, Agrisera), rabbit anti-NBR1 (AS14 2805A, Agrisera), and mouse anti-Flag (A00187, GenScript). Membranes were washed three times with TBS-T and incubated for 1 h at room temperature with alkaline phosphatase-conjugated anti-rabbit (A-3687, Sigma) or anti-mouse (M-9637, Sigma) secondary antibodies diluted 1:1000. Signal was detected using NBT/BCIP substrate solution and imaged using a GelDoc Go Gel Imaging System (Bio-Rad).

Band intensities were quantified using ImageJ software (Schindelin et al., 2012). Images were first processed using the Subtract Background function with a rolling ball radius of 50 pixels. Lanes were selected using rectangular regions encompassing the central third of each lane to minimize edge effects. Densitometric profiles were generated using the Plot Lanes function, and band intensities were calculated as the integrated area above the local background.

Autophagic flux and protein turnover were estimated from densitometric analyses of immunoblots as described by Robert et al. (2025). For GFP-based reporters (35S:GFP–ATG8 and Super:ABI5–GFP), the ratio between free GFP and the corresponding GFP-fusion protein was calculated. For the mCherry–YFP–NBR1 reporter, the ratio between processed mCherry–YFP and full-length mCherry–YFP–NBR1 signals was calculated.

### Co-immunoprecipitation assays

Co-immunoprecipitation assays were performed using transgenic plants expressing ABI5–GFP or ABI5–Flag. Lines use as negatives controls GRX1–roGFP2 (Meyer et al., 2007) and BIM1–Flag (Liang et al., 2018) for GFP- and Flag-based assays were kindly provided by Andreas Meyer (Faculty of Agriculture, University of Bonn) and Georgina Fabro (CIQUIBIC, FCQ-CONICET)), respectively. For ABI5–GFP experiments, leaf tissue from *Pma*-infected plants was collected at 1 day post inoculation (dpi). For ABI5–Flag assays, 14-day-old seedlings grown on 0.5× MS agar medium were used.

Plant tissue was homogenized in extraction buffer containing 50 mM Tris-HCl (pH 7.5), 150 mM NaCl, 10% glycerol, 1 mM EDTA, 0.1% NP-40, 10 mM N-ethylmaleimide (NEM), protease inhibitor cocktail (Roche), and 1 mM PMSF. Extracts were centrifuged at 13,500 rpm for 20 min at 4 °C, and the supernatants were filtered through Miracloth.

For each sample, 10% of the extract was reserved as input and the remaining extract was incubated with Dynabeads Protein A (1001D, Thermo Fischer Scientific) coupled to anti-NBR1 antibodies (AS14 2805A, Agrisera) at 4 °C. ABI5–GFP extracts were incubated overnight, whereas ABI5–Flag extracts were incubated for 4 h. Beads were washed three times with extraction buffer and bound proteins were eluted in 1× SDS-PAGE loading buffer at 70 °C for 10 min. Immunoprecipitated proteins were analyzed by immunoblotting using anti-GFP or anti-Flag antibodies as described above.

### Generation and genotyping of the *nbr1 abi5* double mutant

The *nbr1 abi5* double mutant was generated by crossing homozygous *nbr1* and *abi5-8* plants. F1 progeny derived from the cross, as well as F2 progeny obtained by self-fertilization of F1 plants, were screened on 0.5× MS agar plates supplemented with sulfadiazine (200 μg ml⁻¹) to enrich for individuals carrying the *nbr1* allele. Antibiotic selection for the *abi5-8* allele was not possible because kanamycin resistance had been lost in this line.

Following transplantation to soil, genomic DNA was extracted from leaf tissue of selected F2 individuals and used for PCR genotyping. Homozygous T-DNA insertion mutants were identified using two independent PCR reactions with gene-specific primers and the T-DNA left border primer LBb1.3 (Supplemental Table 1). Plants were scored as homozygous when amplification of the WT allele using gene-specific primers was absent, whereas PCR using a gene-specific primer paired with LBb1.3 yielded a T-DNA-specific fragment of the expected size. PCR amplifications were performed using Pegasus Taq DNA Polymerase (PB-L) or T-Plus DNA Polymerase (Inbio Highway) according to the manufacturers’ instructions under the conditions described in Supplemental Table 2.

### Data analysis and statistics

GraphPad Prism 8 (GraphPad Software) was used for statistical analyses and graph generation. Normality was assessed using the Shapiro–Wilk test, and homogeneity of variances was evaluated using Levene’s test. Depending on the distribution of the data, statistical analyses were performed using one- or two-way analysis of variance (ANOVA) or the Kruskal–Wallis test, as appropriate. Multiple comparisons were corrected using the two-stage linear step-up procedure of Benjamini, Krieger, and Yekutieli. Statistical significance is indicated as follows: P < 0.05 (*), P < 0.01 (**), and P < 0.001 (***).

## RESULTS

### 1. Autophagy and NBR1 are activated upon infection under warming

To investigate whether autophagy is modulated during bacterial infection under elevated temperature, we quantified autophagic flux using the GFP–ATG8a cleavage assay (Thompson et al., 2005). In this system, GFP–ATG8a is delivered to the vacuole during autophagy, where ATG8a is degraded while free GFP accumulates due to its relative stability. Thus, autophagic flux can be estimated as the ratio of free GFP to GFP–ATG8a. Five-week-old Arabidopsis plants expressing GFP–ATG8a were infiltrated with mock or *Pseudomonas cannabina* pv*. alisalensis* (*Pma*, formerly *P. syringae* pv. *maculicola* ES4326; Baltrus et al., (2011); Sarris et al., (2013)) at 5 × 10 CFU ml⁻¹ and incubated at either 22 °C or 29 °C, the latter used as a warming condition. Autophagic flux was not significantly altered by temperature in plants infiltrated with mock (Figure 1a). In contrast, *Pma* infection led to an increase in autophagic flux at 29 °C compared to 22 °C at 2 days post-inoculation (dpi). Consistently, *Pma*-inoculated seedlings transferred to 29 °C showed increased autophagic flux and more GFP–ATG8a-labeled puncta, particularly in shoot tissues (Supplemental Figure 1), indicating that warming enhances infection-induced autophagy in both adult plants and seedlings. To determine whether this response is conserved in another commonly used bacterial strain and whether it depends on effector delivery, we analyzed plants infected with *P. syringae* pv. *tomato* DC3000 (*Pst*) and the type III secretion-deficient mutant *Pst-*Δ*hrcC*. Both strains induced a significant increase in autophagic flux at 29 °C (Supplemental Figure 2). This indicates that temperature-enhanced autophagy does not require type III effector delivery and is consistent with activation by PAMP recognition and PTI-associated signaling.

**Figure 1.**
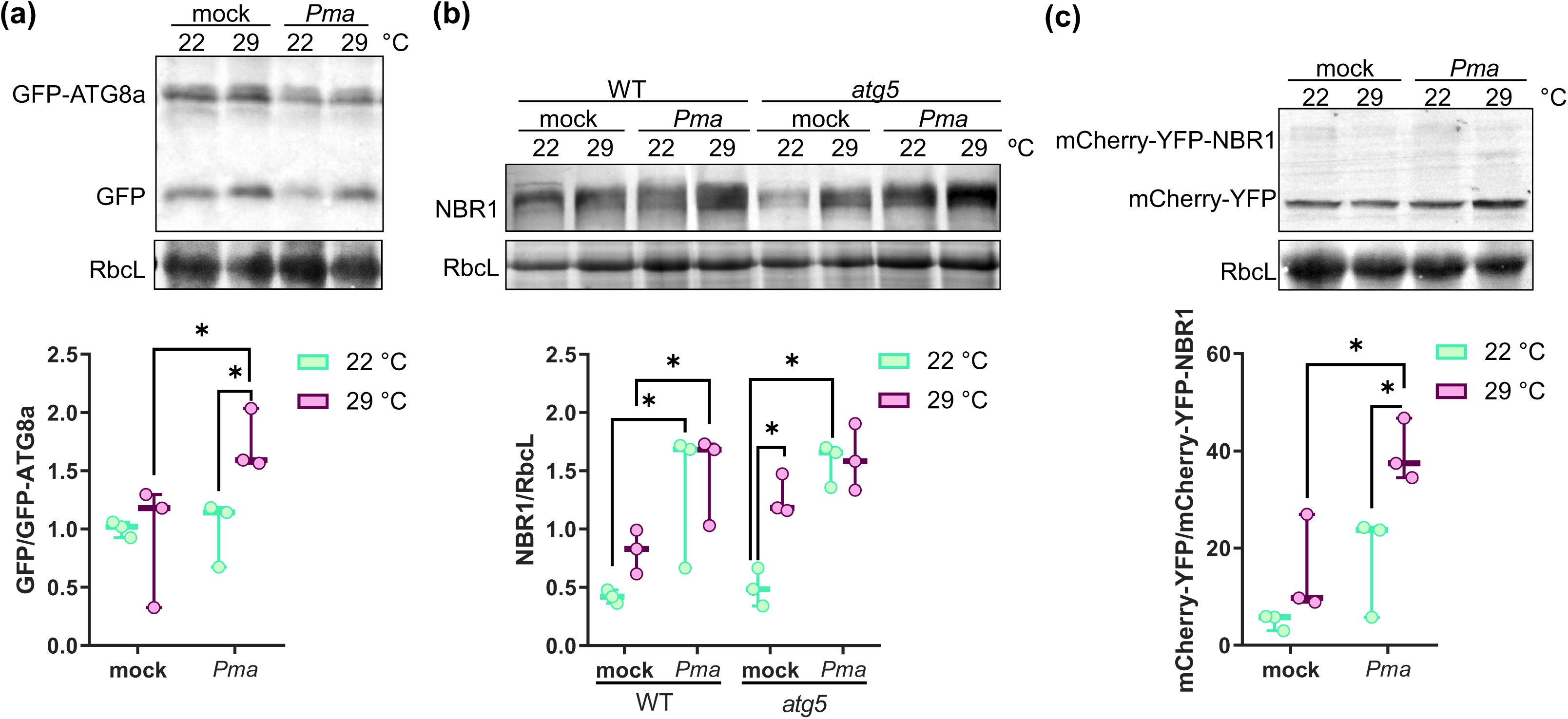
*Pma* infection under warming activates autophagic flux and promotes NBR1 turnover in Arabidopsis. Plants were infiltrated with mock solution or *Pma* at 5 × 10⁵ CFU ml⁻¹, and subsequently incubated at 22 °C or 29 °C. **(a)** Autophagic flux was monitored in GFP–ATG8a plants at 2 days post-inoculation (dpi) as the ratio of free GFP to GFP–ATG8a. **(b)** Immunoblot analysis of endogenous NBR1 protein levels in WT and *atg5* plants at 1 dpi. **(c)** Immunoblot analysis of mCherry–YFP and mCherry–YFP–NBR1 fusion proteins at 1 dpi using an anti-mCherry antibody to assess NBR1 turnover. Rubisco large subunit (RbcL) was used as a loading control. Band intensities were quantified by densitometric analysis, and relative values normalized to the fusion proteins ((a), (c)) or to RbcL ((b)) are shown below each blot. In (a), values are expressed relative to the 22 °C mock condition. Each circle represents an individual plant. Boxes indicate the median and interquartile range, and whiskers represent the minimum and maximum values. Asterisks indicate statistically significant differences (*P < 0.033; two-way ANOVA for (a) and (c), or Kruskal–Wallis test for (b), followed by false discovery rate correction using the Benjamini, Krieger and Yekutieli two-stage linear step-up procedure).

We next assessed the levels of the selective autophagy receptor NBR1, a well-established autophagy receptor that accumulates when autophagic degradation is impaired (Üstün et al., 2018). Immunoblot analysis revealed that NBR1 protein levels increased upon infection, particularly under warming conditions (Figure 1b). Importantly, NBR1 accumulation was strongly enhanced in *atg5* mutants, which are defective in autophagosome formation (Thompson et al., 2005), indicating that NBR1 turnover depends on a functional autophagy pathway.

To directly monitor NBR1 turnover, we used plants expressing the tandem fluorescent reporter mCherry–YFP–NBR1 (Svenning et al., 2011). Upon autophagic degradation, the fusion protein is processed, leading to the release of free mCherry–YFP, which can be detected by immunoblotting. Thus, NBR1 turnover was estimated as the ratio of free mCherry–YFP to mCherry–YFP–NBR1. Under infection and warming conditions, we observed an increase in the free mCherry–YFP/mCherry–YFP–NBR1 ratio compared to mock or single treatments, consistent with enhanced NBR1 turnover (Figure 1c). Taken together, these results strongly suggest that autophagy is activated during bacterial infection under warming conditions. Furthermore, autophagy-dependent turnover of NBR1 increased during infection and was further enhanced under elevated temperatures.

### 2. NBR1-mediated autophagy contributes to resistance under warming

To assess the functional relevance of NBR1-mediated autophagy under warming conditions, we quantified susceptibility to *Pma* bacterial infection in WT, *nbr1*, and *atg5* plants, as well as in plants overexpressing NBR1 (35S:mCherry–YFP–NBR1), at 22 °C or 29 °C (Figure 2). No significant differences in bacterial growth were observed among genotypes at 22 °C (Figure 2a). In contrast, all genotypes showed increased bacterial proliferation at 29 °C, indicating that elevated temperature generally enhances susceptibility to infection as previously reported by Huot et al. (2017). Notably, the magnitude of this increase differed markedly among genotypes. Autophagy mutants *nbr1* and *atg5* showed higher bacterial titers compared to WT under warming conditions (Figure 2a, Supplemental Figure 3a). To compare the temperature-dependent increase in susceptibility across genotypes, we calculated a susceptibility index (SI) by normalizing bacterial growth at 29 °C to that at 22 °C for each genotype (Figure 2b). While the SI in WT plants was ∼5, *nbr1* and *atg5* mutants displayed SI values of 100-130, indicating a markedly stronger temperature-dependent increase in bacterial growth (Figure 2b, Supplemental Figure 3b). Notably, the increased susceptibility observed in *atg5* mutants is consistent with impaired autophagic flux and defective turnover of NBR1 (Figure 1b), supporting a role for autophagy-dependent processes in this response. In contrast, plants overexpressing NBR1 did not show a significant increase in bacterial growth at 29 °C relative to 22 °C and displayed an SI of 0.82, lower than that of WT (Figure 2a, b), indicating enhanced tolerance under warming conditions. Moreover, a similar susceptibility pattern was observed upon infection with *P. syringae* pv. *tomato* DC3000 (*Pst*, Supplemental Figure 4), indicating that the enhanced temperature-dependent susceptibility of *nbr1* is not restricted to *Pma*. In addition to bacterial growth, chlorophyll content was measured as a physiological indicator of disease impact (Supplemental Figure 3c). Although differences in chlorophyll levels were relatively modest, *nbr1* plants consistently maintained higher chlorophyll contents than WT plants, supporting the possibility that chlorophyll catabolism is altered in the mutant.

**Figure 2.**
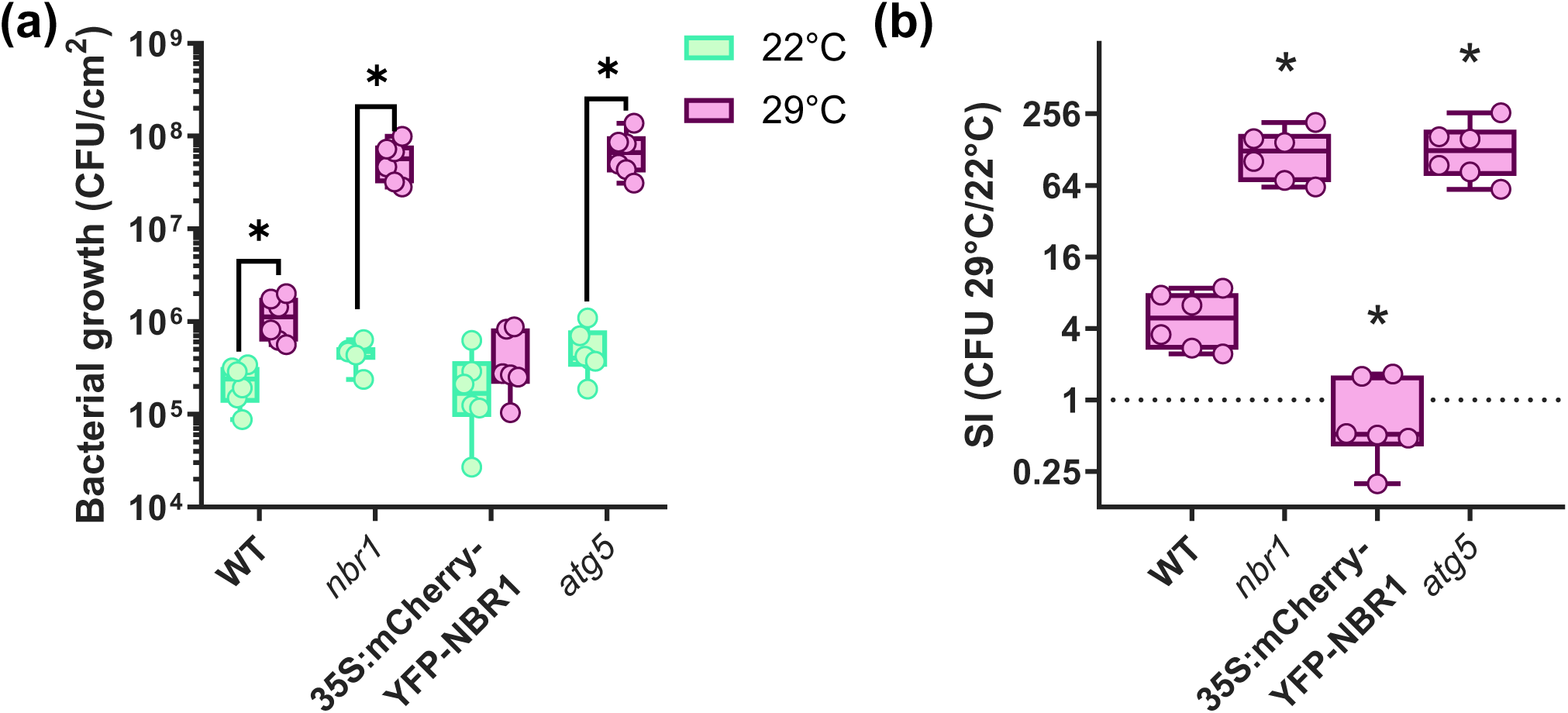
NBR1-mediated selective autophagy limits bacterial growth under warming conditions. WT plants, mutants impaired in NBR1 (*nbr1*) or core autophagy (*atg5*), and plants overexpressing NBR1 (35S:mCherry–YFP–NBR1) were infiltrated with *Pma* at 5 × 10⁵ CFU ml⁻¹, and subsequently incubated at 22 °C or 29 °C. Bacterial growth was quantified at 2 days post-inoculation. **(a)** Bacterial titers expressed as CFU per cm² of leaf tissue. **(b)** Susceptibility index (SI), calculated as the ratio between bacterial titers at 29 °C and the mean bacterial titer at 22 °C for the same genotype. These results were reproducible across three independent experiments (Supplemental Figure 3). Each circle represents an individual plant. Boxes indicate the median and interquartile range, and whiskers represent the minimum and maximum values. Asterisks indicate statistically significant differences (*P < 0.033; Kruskal–Wallis test followed by false discovery rate correction using the Benjamini, Krieger and Yekutieli two-stage linear step-up procedure). In (b), statistical comparisons were performed relative to WT.

To determine whether the contribution of NBR1 to defense under elevated temperature depends on type III secretion or on effector-triggered immunity (ETI), we compared bacterial growth and SI in plants infected with virulent *Pst*, the type III secretion (T3SS)-deficient mutant *Pst-*Δ*hrcC* and an AvrRpt2-expressing strain (*Pst-AvrRpt2*). While *nbr1* plants displayed increased susceptibility to *Pst*, no significant differences between WT and *nbr1* plants were observed upon infection with *Pst-*Δ*hrcC* or *Pst-AvrRpt2* (Supplemental Figure 4). These results indicate that the NBR1-dependent contribution to resistance under warming is most evident during infection with a virulent strain carrying a functional T3SS, rather than during the response to a secretion-deficient strain. This is consistent with previous evidence showing that *Pst* activates autophagy in a T3SS-dependent manner and that NBR1-mediated selective autophagy can counteract effector-associated disease progression (Üstün et al., 2018). Thus, NBR1, or NBR1-dependent downstream processes, may limit susceptibility promoted by one or more Pseudomonas effectors under elevated temperature. By contrast, the absence of a detectable *nbr1* phenotype during AvrRpt2-triggered immunity may reflect the strength of this ETI response, which could override or mask the NBR1-dependent contribution observed during virulent infection. Taken together, these findings indicate that NBR1-mediated selective autophagy helps restrict bacterial proliferation under warming conditions, particularly in the context of effector-mediated susceptibility.

### 3. NBR1 negatively regulates ABA signaling under warming

Because phytohormone signaling plays a central role in plant immunity and is strongly influenced by temperature, we next examined whether NBR1 affects hormone pathways associated with bacterial infection under warming. Elevated temperature has been reported to enhance disease susceptibility in Arabidopsis, in part through suppression of salicylic acid (SA)-mediated defenses and modulation of other hormonal pathways (Huot et al., 2017). Among these, auxin signaling has been linked to increased susceptibility to bacterial pathogens and antagonism of SA-dependent defenses (Kazan and Manners, 2009). We therefore quantified SA and indole-3-acetic acid (IAA) levels to investigate whether altered hormonal responses contribute to the enhanced susceptibility of *nbr1* plants under warm conditions. IAA content showed a slight increase at 29 °C compared to 22 °C, but no significant differences were observed between genotypes or infection conditions (Figure 3a). In contrast, *Pma* infection strongly induced SA accumulation and the expression of SA-responsive genes, including *PR1* and *ICS1* (Figure 3b, S5a–b). However, both SA levels and *PR1* expression were reduced under warming conditions, consistent with previous reports. Importantly, these responses were comparable between WT and *nbr1* plants, indicating that NBR1 deficiency does not substantially alter SA accumulation or SA-responsive gene expression under these conditions. We also quantified jasmonic acid (JA), since this hormone might contribute to immune responses against hemibiotrophic bacterial pathogens (Ellis et al., 2002; Zhang et al., 2017). Although *Pma* infection induced JA accumulation in WT plants at 22 °C, this response was impaired in *nbr1* plants and was largely suppressed by elevated temperature in both genotypes (Figure 3c). Thus, while NBR1 appears to contribute to infection-induced JA accumulation under control temperature, this effect was not maintained under warming conditions, where JA levels remained low. Therefore, neither SA nor JA responses appeared to fully explain the enhanced susceptibility of *nbr1* plants specifically under elevated temperature.

**Figure 3.**
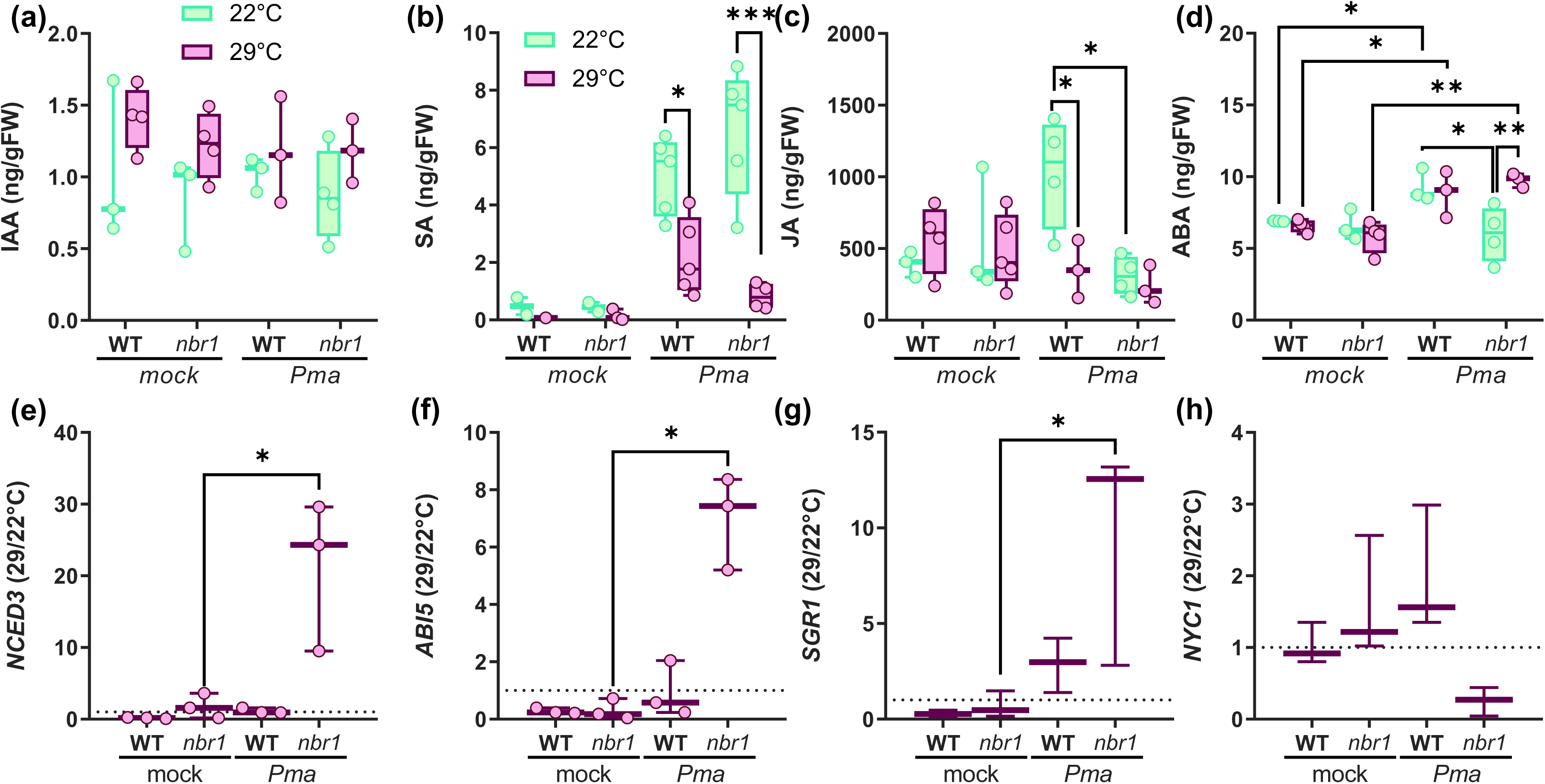
NBR1 negatively regulates ABA accumulation and ABA-associated signaling during bacterial infection under warming. WT and *nbr1* plants were infiltrated with mock solution or *Pma* at 5 × 10⁵ CFU ml⁻¹, and subsequently incubated at 22 °C or 29 °C. **(a-d)** Indole-3-acetic acid (IAA), salicylic acid (SA), jasmonic acid (JA) and abscisic acid (ABA) concentrations measured at 2 days post-inoculation. **(e-h)** Temperature sensitivity index for ABA biosynthesis and signaling–associated genes *NCED3*, *ABI5*, *SGR1* and *NYC1*, calculated as the ratio between transcript levels at 29 °C and the mean transcript levels at 22 °C for the same genotype and treatment. A dotted horizontal line at y = 1 indicates no temperature-dependent change in transcript abundance. Data represent the mean ± SEM of 3–4 biological replicates; circles indicate individual replicates. Asterisks indicate statistically significant differences (\**P* < 0.05, \*\**P* < 0.002; two-way ANOVA or Kruskal–Wallis test followed by false discovery rate correction using the Benjamini, Krieger and Yekutieli two-stage linear step-up procedure).

In contrast, ABA accumulation and signaling showed a genotype-specific response that was most evident during infection under warming conditions. In WT plants, *Pma* infection induced a moderate increase in ABA levels at both temperatures (Figure 3d). In *nbr1* plants, however, infection did not significantly alter ABA levels at 22 °C, whereas it caused a pronounced increase under warm conditions. These results indicate that NBR1 deficiency specifically enhances ABA accumulation during bacterial infection at elevated temperature, prompting us to focus on this pathway. To further characterize this response, we analyzed the expression of ABA-related genes. *NCED3*, a key enzyme in ABA biosynthesis, was not significantly affected by temperature in WT plants, but showed increased expression in *nbr1* plants at 29 °C (Supplemental Figure 5c). Similarly, the ABA-responsive transcription factor *ABI5* and its downstream target *SGR1* showed elevated expression in *nbr1* plants compared to WT upon infection under warm conditions (Supplemental Figure 5d-e). In contrast, expression of the ABI5 target gene *NYC1* was reduced in *nbr1* plants at 29 °C (Supplemental Figure 5f), suggesting differential regulation of ABA-responsive downstream genes.

To quantify the effect of warming on ABA-associated transcriptional responses, we calculated a temperature sensitivity index, defined as the ratio between transcript levels at 29 °C and those at 22 °C (Figure 3d-f). While this ratio was not significantly altered in WT plants upon infection, *NCED3*, *ABI5* and *SGR1*, but not *NYC1*, displayed a markedly higher temperature responsiveness in infected *nbr1* plants, indicating an exacerbated ABA-associated transcriptional response under warming. Taken together, these results indicate that, under our conditions, SA accumulation and signaling are strongly suppressed by elevated temperature but are not substantially affected by NBR1 deficiency. JA induction depends on NBR1 at 22 °C but is not sustained under warming. By contrast, ABA accumulation and ABA-associated gene expression are specifically exacerbated in infected *nbr1* plants at elevated temperature, supporting a major role for ABA in the warming-dependent susceptibility of *nbr1*.

### 4. NBR1 interacts with ABI5 and regulates its turnover

Given the central role of ABI5 in ABA signaling and the enhanced ABA responses observed in *nbr1* plants under warm conditions, we investigated whether NBR1 directly regulates ABI5. Previous work has shown that NBR1 can interact with ABA-related transcription factors, including ABI5, suggesting that ABI5 could be a target of selective autophagy (Tarnowski et al., 2020). First, to assess how temperature and infection affect ABI5 protein accumulation and whether this is regulated by NBR1, we generated WT, *nbr1* and *atg5* plants expressing Flag-C-term-tagged ABI5 under the control of the constitutive pSuper promoter (Qi et al., 2020). Immunoblot analysis confirmed the absence of NBR1 in the *nbr1* background (Figure 4a). In WT plants, NBR1 protein levels increased upon infection and under warming conditions, consistent with the accumulation patterns observed previously (Figure 1b). ABI5–Flag protein was detected at low levels under mock conditions, with slightly higher accumulation in *nbr1* compared to WT. Upon *Pma* infection, ABI5–Flag levels increased in both genotypes, but this induction was markedly stronger in *nbr1* plants. Interestingly, although warming also promoted ABI5 accumulation, ABI5–Flag levels in infected plants were lower at 29 °C compared to 22 °C, suggesting a temperature-dependent modulation of ABI5 stability (Figure 4a).

**Figure 4.**
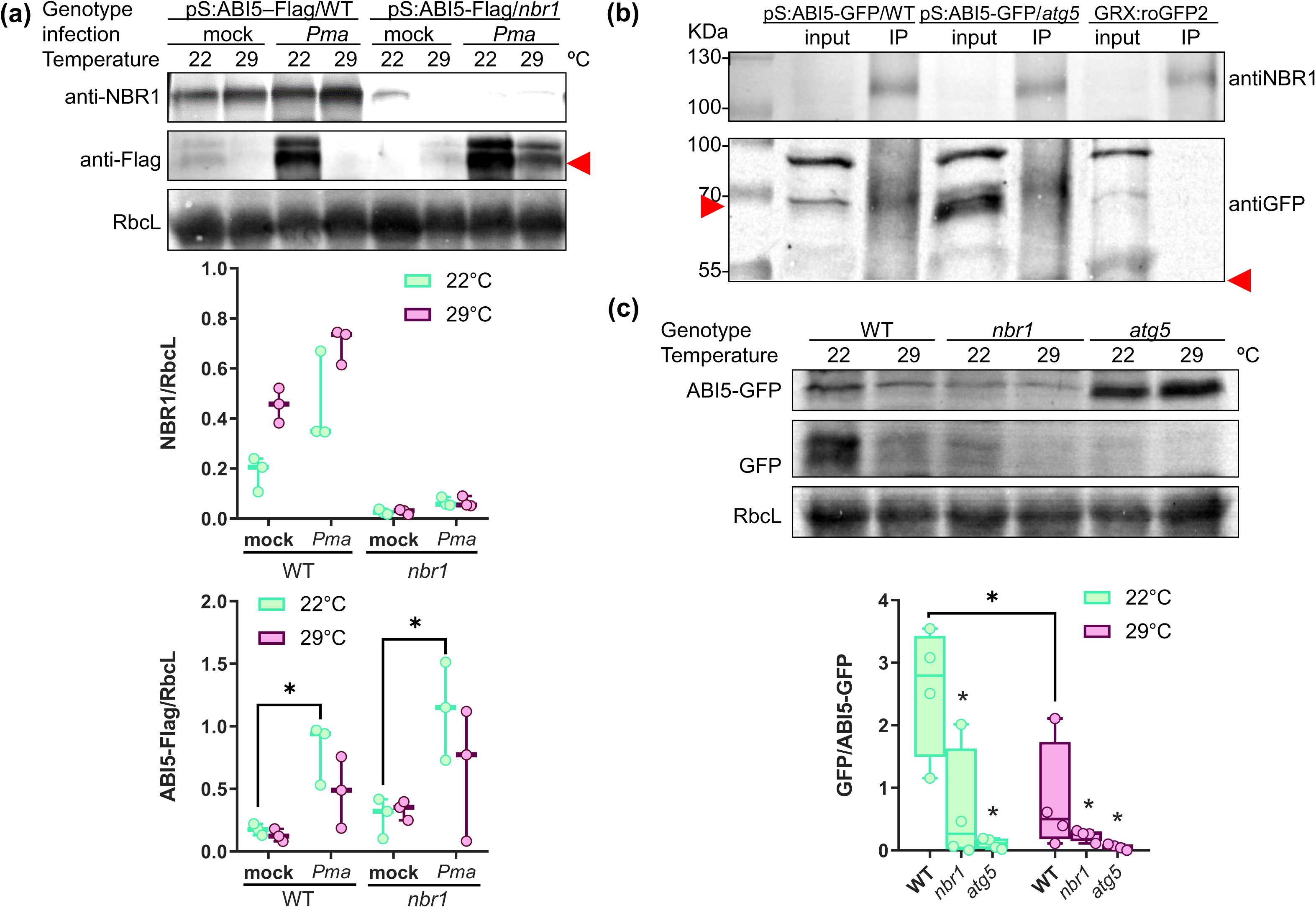
NBR1 interacts with ABI5 and regulates its autophagy-dependent turnover during bacterial infection. Plants expressing pSuper:ABI5-Flag or pSuper:ABI5-GFP in WT, *nbr1*, or *atg5* backgrounds were infiltrated with mock solution or *Pma* at 5 × 10⁵ CFU ml⁻¹, and incubated at 22 °C or 29 °C for 1 day. **(a)** Representative immunoblots are shown, and densitometric quantification of NBR1 and ABI5-Flag protein levels relative to the Rubisco large subunit (RbcL) is shown below blots. **(b)** Co-immunoprecipitation of ABI5-GFP using anti-NBR1 magnetic beads in *Pma*-infected pSuper:ABI5-GFP/WT and pSuper:ABI5-GFP/*atg5* plants at 22 °C. Input and immunoprecipitated fractions are shown. GRX–roGFP2 plants were used as a negative control. In (a) and (b), red arrows indicate the positions of tagged proteins. A non-specific band migrating between ∼70 and 100 kDa was detected in input samples. Ponceau staining of RbcL is shown as a loading control. **(c)** Immunoblot analysis of ABI5-GFP turnover in *Pma*-infected pSuper:ABI5-GFP plants in WT, *nbr1*, and *atg5* backgrounds. Autophagy-dependent turnover was quantified by densitometric analysis as the ratio of free GFP to ABI5-GFP. Representative immunoblots and corresponding quantification analysis are shown. Each circle represents an individual plant. Boxes indicate the median and interquartile range, and whiskers represent the minimum and maximum values. Statistical analysis was performed using two-way ANOVA on log2-transformed ratio values followed by false discovery rate correction using the Benjamini, Krieger and Yekutieli two-stage linear step-up procedure. Asterisks indicate statistically significant differences (*P < 0.05) relative to WT or between treatments, as indicated.

To test whether NBR1 physically associates with ABI5 *in planta*, we performed co-immunoprecipitation assays using transgenic lines expressing ABI5 fused to either GFP or Flag tags (Qi et al., 2020). Consistent with that reported by Tarnowski et al. (2020), following immunoprecipitation with anti-NBR1 magnetic beads, ABI5-GFP and ABI5-Flag, but not tagged control proteins, were specifically detected in immunoprecipitated fractions, indicating that NBR1 and ABI5 associate *in planta* (Figure 4b, Supplemental Figure 6). To determine whether ABI5 is degraded through autophagy, we analyzed ABI5 turnover using transgenic lines expressing ABI5–GFP in WT, *nbr1*, and *atg5* backgrounds. Similar to established GFP-based autophagy reporters (Thompson et al., 2005), cleavage of the fusion protein releases free GFP, allowing estimation of protein turnover by immunoblotting as the ratio of free GFP to full-length ABI5–GFP. Under mock conditions, ABI5 turnover was reduced in both *nbr1* and *atg5* plants compared to WT, with the strongest effect observed at 29 °C, where both mutants exhibited similarly low turnover rates (Supplemental Figure 7). These results indicate that basal ABI5 degradation depends on autophagy and suggest that NBR1-mediated selective autophagy becomes a major contributor to ABI5 turnover under warm conditions. Upon infection, WT plants showed a modest reduction in ABI5 turnover at 29 °C relative to 22 °C (Figure 4c). In contrast, both *nbr1* and *atg5* backgrounds displayed impaired turnover, with the GFP/ABI5–GFP ratio remaining significantly lower than in WT. While *nbr1* plants exhibited only a partial reduction in ABI5 turnover at 22 °C, the turnover defect at 29 °C was comparable to that observed in *atg5* mutants, indicating that NBR1-mediated selective autophagy becomes a major determinant of ABI5 turnover under warm conditions. Taken together, these results identify ABI5 as an autophagy-regulated protein whose turnover is promoted during infection under warming conditions. These findings further suggest that the contribution of NBR1 to ABI5 degradation increases at elevated temperature, revealing a temperature-dependent role for NBR1-mediated selective autophagy in the regulation of ABA signaling.

### 5. ABI5 contributes to NBR1-dependent susceptibility under warming

To determine whether the loss of NBR1-mediated ABI5 degradation contributes to the increased susceptibility of *nbr1* plants under warm conditions, we generated *nbr1 abi5* double mutants (Supplemental Figure 8). WT, *nbr1*, *abi5*, and *nbr1 abi5* plants were grown at 22 °C and infected with *Pma* and incubated at 22 °C or 29 °C for 2 days prior to bacterial quantification (Figure 5). Consistent with previous results, WT plants displayed increased bacterial growth under warm conditions, whereas *nbr1* plants exhibited a markedly stronger increase in bacterial proliferation and SI (Figure 5). In contrast, *abi5* mutants showed only a modest and non-significant increase in bacterial growth at 29 °C. Importantly, loss of ABI5 not only suppressed the enhanced susceptibility of *nbr1* plants but reduced bacterial growth and SI values to levels even lower than those observed in WT plants (Figure 5). These results indicate that the presence of ABI5 is needed for the increased susceptibility observed in *nbr1* plants under warming conditions.

**Figure 5.**
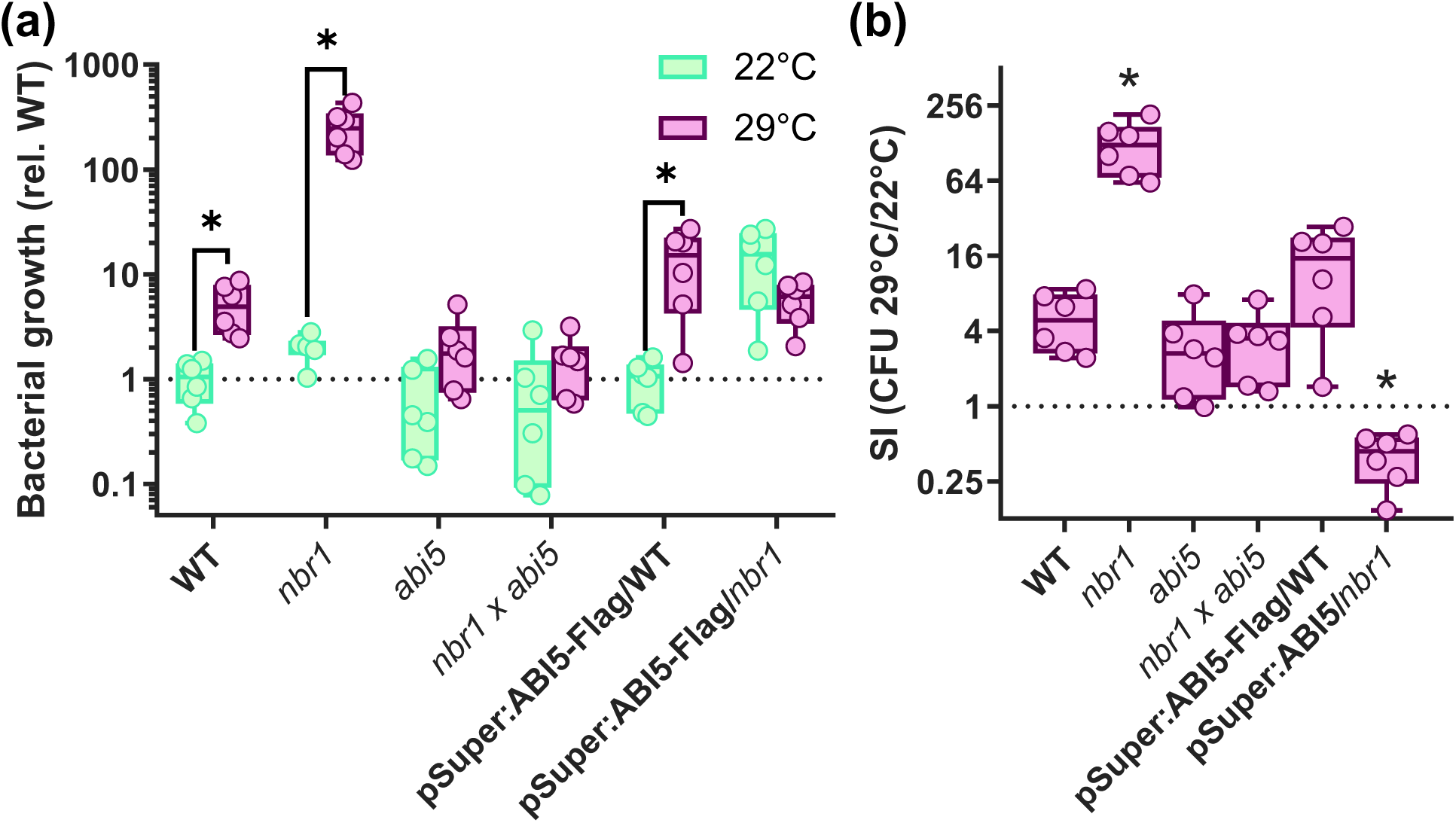
ABI5 contributes to NBR1-dependent susceptibility under warming conditions. WT, *nbr1*, *abi5*, *nbr1 abi5*, pSuper:ABI5–3×Flag/WT and pSuper:ABI5–3×Flag/*nbr1* plants were infiltrated with *Pma* at 5 × 10⁵ CFU ml⁻¹ and subsequently incubated at 22 °C or 29 °C. **(a)** Bacterial growth was quantified at 2 days post-inoculation (dpi) and expressed relative to WT plants grown at 22 °C, which were assigned a value of 1. A dotted horizontal line indicates this reference value. **(b)** Susceptibility index (SI), calculated as the ratio between bacterial titers at 29 °C and the mean bacterial titer at 22 °C for the same genotype. A dotted horizontal line at y = 1 indicates equal bacterial growth at 22 °C and 29 °C (i.e. no temperature-dependent effect on susceptibility). Each circle represents an individual plant. Boxes indicate the median and interquartile range, and whiskers represent the minimum and maximum values. Asterisks indicate statistically significant differences (*P < 0.05) as determined by Kruskal–Wallis test followed by false discovery rate correction using the Benjamini, Krieger and Yekutieli two-stage linear step-up procedure. In (b), statistical comparisons were performed relative to WT.

To further evaluate the contribution of ABI5, we analyzed transgenic lines expressing ABI5-Flag under the constitutive pSuper promoter that were previously used for Western blot experiments (Figure 4a, Supplemental Figure 6). While pSuper:ABI5–3×Flag/WT plants displayed bacterial growth levels similar to WT at 22 °C, they exhibited a stronger increase in bacterial proliferation under warming conditions, resulting in elevated SI values (Figure 5). These results are consistent with increased ABI5 activity promoting susceptibility during infection under elevated temperature. In contrast, pSuper:ABI5–3×Flag/*nbr1* plants displayed elevated bacterial growth already at 22 °C and showed little additional increase at 29 °C. Accordingly, their SI values remained lower than those observed in pSuper:ABI5–3×Flag/WT plants (Figure 5). This suggests that the absence of NBR1 sensitizes plants to ABI5-dependent responses, leading to a constitutively susceptible state and leaving little capacity for further disease enhancement at elevated temperature.

Taken together, these results demonstrate that ABI5 contributes to temperature-enhanced susceptibility during bacterial infection and support the conclusion that NBR1 restricts susceptibility under warming conditions, at least in part, through negative regulation of ABI5.

## DISCUSSION

In this study, we show that NBR1-mediated selective autophagy contributes to limiting bacterial susceptibility in Arabidopsis particularly under elevated temperature conditions, through the negative regulation of ABA signaling. This temperature dependence is a central feature of our findings, as the contribution of NBR1 to resistance became most evident under warming, a condition known to favor bacterial disease development and to reconfigure plant immune and hormonal responses (Huot et al., 2017). Elevated temperature has been associated with increased ABA responses during bacterial infection, often coinciding with enhanced disease susceptibility (De Torres-Zabala et al., 2007; Cao et al., 2011; Mine et al., 2017). Consistent with this view, the enhanced susceptibility of *nbr1* plants under warming was accompanied by a stronger activation of ABA-associated responses, whereas transcriptional markers of SA signaling remained largely unaffected despite the reduced SA accumulation observed in *nbr1* plants (Supplemental Figure 5a-b, Figure 3b). These observations suggest that, under elevated temperature, the increased susceptibility of *nbr1* plants is more closely associated with altered ABA signaling than with a major impairment in SA-dependent defenses. NBR1 also influenced JA accumulation during infection. Whereas WT plants accumulated JA following infection at 22 °C, this response was largely absent in *nbr1* plants and was suppressed by warming in both genotypes (Figure 3c). Thus, elevated temperature appears to shift the hormonal balance during infection by enhancing ABA-associated responses while attenuating JA accumulation, generating a physiological context that may favor bacterial susceptibility and increase the relevance of NBR1-mediated regulation.

Bacterial infection under warming conditions enhanced autophagic flux and promoted NBR1 turnover (Figure 1, Supplemental Figure 1, S2), indicating activation of selective autophagy under conditions in which ABA signaling becomes particularly relevant for disease outcome. More broadly, our results indicate that autophagy contributes to limiting bacterial susceptibility under warming, as both *atg5* and *nbr1* mutants displayed enhanced disease phenotypes (Figure 2). Recent studies have begun to reveal that autophagy regulates ABA signaling through the selective turnover of multiple pathway components. For example, ABA receptors such as PYR1, PYL3 and PYL4 accumulate in autophagy-deficient mutants, while the ABA-responsive transcription factors bZIP67 and ABI5 are stabilized in *atg* backgrounds (Contreras et al., 2025; Zhu et al., 2026). In addition, NBR1 was previously reported to associate with ABI3, ABI4 and ABI5 *in planta* (Tarnowski et al., 2020), although whether these transcription factors are selectively degraded through NBR1-mediated autophagy has remained unresolved. Notably, ABI5 interaction requires full-length NBR1, whereas ABI3 and ABI4 can associate with NBR1 variants lacking the UBA domain, suggesting that distinct recognition mechanisms may operate for different ABA-responsive transcription factors (Tarnowski et al., 2020). Our findings provide direct support for this idea by showing that ABI5 accumulates in both *nbr1* and *atg5* mutants and that its turnover is partially dependent on NBR1-mediated autophagy (Figure 4). The stronger impairment of ABI5 turnover in *atg5* relative to *nbr1* plants at 22 °C, together with the similar defects observed in both mutants at 29 °C (Figure 4c, Supplemental Figure 7), suggests that NBR1 becomes a major contributor to ABI5 degradation under warm conditions, while additional autophagy-dependent mechanisms may participate under control temperatures. More recently, AtATG8H was shown to interact specifically with ABI3 and ABI5, further supporting the idea that ABA-responsive transcription factors can be directly recognized by the autophagy machinery (Contreras et al., 2025). Together with our findings, these observations suggest that autophagy regulates ABA signaling at multiple levels, including hormone perception and transcriptional output. Such multilayered control may provide the plasticity required to fine-tune ABA-dependent responses under different environmental and developmental conditions. Notably, our results indicate that the contribution of NBR1 to ABI5 turnover increases at elevated temperature (Figure 4b, S7), suggesting that warming enhances the dependence of ABA signaling on selective autophagy-mediated regulation.

The genetic suppression of the *nbr1* susceptibility phenotype by *abi5* mutation identifies ABI5 as a key downstream component of NBR1-mediated defense under warm conditions (Figure 5). ABI5 is a central transcriptional regulator of ABA signaling and controls the expression of numerous stress-responsive genes (Skubacz et al., 2016). Although ABI5 has been extensively studied in seed maturation and abiotic stress responses, its role in plant–pathogen interactions remain comparatively less explored. Notably, *abi5* mutants display enhanced resistance to *P. syringae* expressing the type III effector HopAM1 (Goel et al., 2008). Because HopAM1 promotes activation of ABA-responsive pathways, these findings established an early connection between bacterial effector activity, ABA signaling, ABI5 function and bacterial susceptibility. Additional support for effector-mediated manipulation of ABA signaling comes from AvrPtoB, a *Pst* effector reported to target the ABA 8′-hydroxylases CYP707A1 and CYP707A3, thereby reducing ABA catabolism and promoting ABA accumulation and virulence (Liu et al., 2022). Thus, although a direct Pseudomonas effector targeting ABI5 has not, to our knowledge, been reported, effector-mediated activation of ABA signaling may increase the dependence on NBR1-dependent control of ABI5 homeostasis. Our results extend this connection by showing that the contribution of ABI5 to bacterial susceptibility becomes particularly evident under elevated temperature (Figure 5).

The downstream mechanisms by which ABI5 promotes bacterial susceptibility under warming remain unclear. Our data suggests that altered chloroplast-associated processes could contribute to this phenotype. In infected *nbr1* plants, increased *SGR1* expression was not accompanied by induction of *NYC1*, and *nbr1* plants retained higher chlorophyll content than WT plants (Figure 3g–h, Supplemental Figure 5e-f, Supplemental Figure 3c). Because chlorophyll catabolites can influence immune responses and hypersensitive cell death during Pseudomonas infection (Mur et al., 2010), altered chlorophyll metabolism may represent one way by which enhanced ABA/ABI5 signaling affects disease outcome. In addition, since ABA biosynthesis is initiated in plastids and NBR1 has been implicated in chloroplast quality control under stress conditions (Lee et al., 2023), plastid homeostasis may provide another link between selective autophagy, ABA signaling and susceptibility under warming. However, further studies will be required to determine whether these alterations contribute directly to the susceptibility phenotype observed in *nbr1* plants.

Consistent with the idea that NBR1 becomes particularly important when bacterial virulence strategies operate under elevated temperature, the enhanced susceptibility of *nbr1* was observed under warming during infection with virulent Pseudomonas strains but was largely absent following infection with either the T3SS mutant *Pst-*Δ*hrcC* or the ETI-inducing strain *Pst-AvrRpt2* (Figure 2, Supplemental Figure 4). This pattern suggests that the NBR1-dependent contribution to resistance becomes particularly relevant under elevated temperature during compatible interactions with a functional T3SS, rather than during responses to a secretion-deficient strain or a strong AvrRpt2-triggered ETI response. Warming may therefore create a host physiological state in which effector-mediated susceptibility, including effector-driven ABA pathway manipulation, is more effective or has stronger consequences for disease outcome. In this regard, it is noteworthy that elevated temperature has been reported to enhance the translocation of bacterial effector proteins into plant cells (Huot et al., 2017), and that *Pst* activates autophagy in a T3SS effector-dependent manner (Üstün et al., 2018). Moreover, NBR1-dependent selective autophagy has been shown to counteract HopM1-mediated disease development and water-soaked lesion formation during Pseudomonas infection. Thus, NBR1-mediated regulation of ABA/ABI5 signaling may act as a protective layer that limits effector-associated susceptibility specifically under environmental conditions that favor pathogen virulence. The absence of a detectable *nbr1* phenotype during *AvrRpt2*-triggered immunity may reflect the strength of this ETI response, which could override or mask the contribution of NBR1-dependent selective autophagy observed during virulent infection under warming (Supplemental Figure 3, Fig.1, Supplemental Figure 2).

Together, our findings identify NBR1-mediated selective autophagy as a mechanism that restrains ABA/ABI5-dependent susceptibility during bacterial infection under elevated temperature. By promoting ABI5 turnover, NBR1-mediated selective autophagy may function as a temperature-dependent buffering mechanism that prevents excessive ABA signaling when warm conditions favor bacterial virulence (Figure 6). This work links selective autophagy to the hormonal reprogramming of plant immunity under high temperature and suggests that NBR1-dependent control of ABA signaling may be an important component of disease resistance in warming environments.

**Figure 6.**
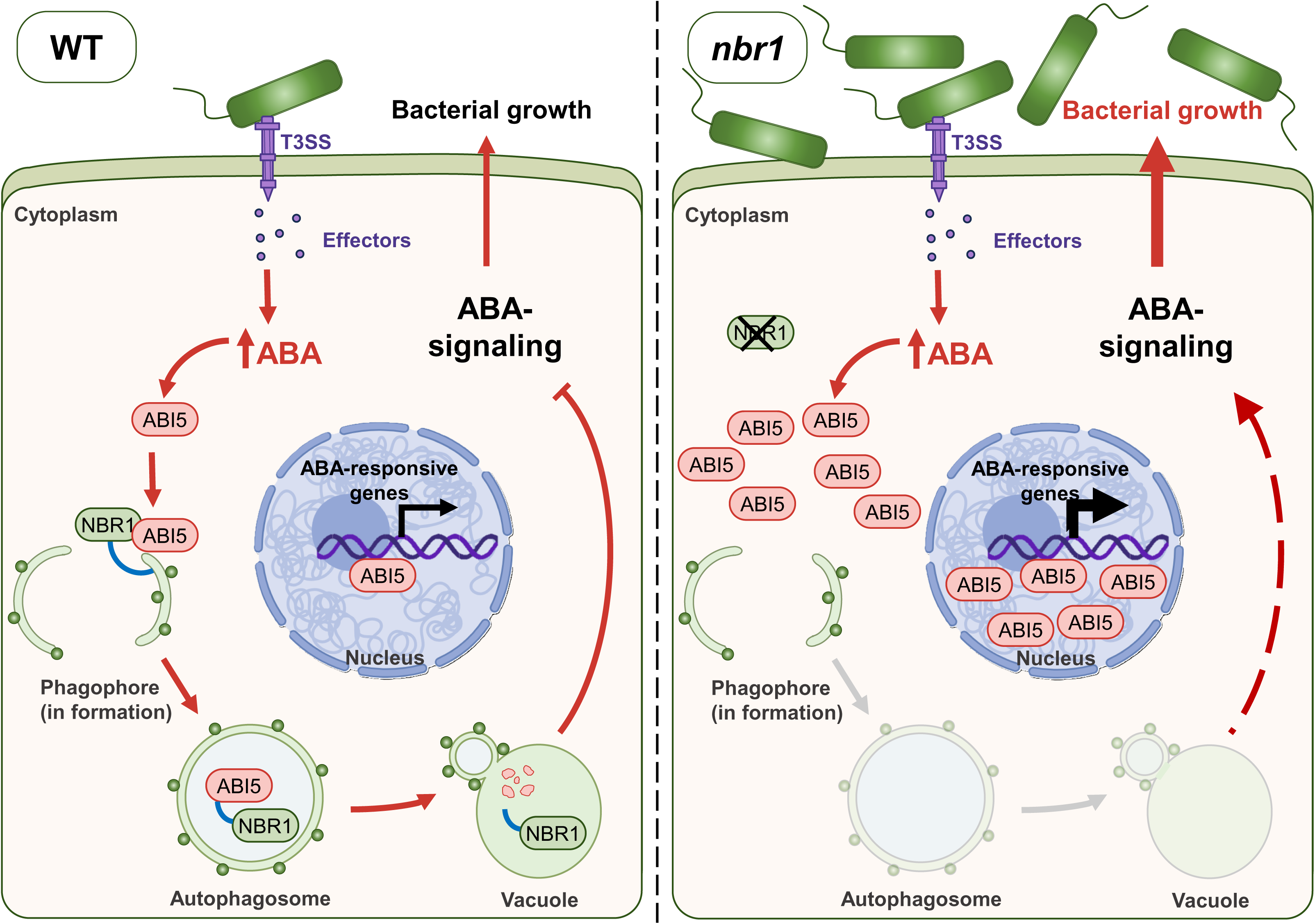
Proposed model for the role of NBR1-mediated selective autophagy in limiting ABA-dependent susceptibility under warming conditions. During bacterial infection under elevated temperature, bacterial effectors promote ABA accumulation and ABA signaling, contributing to susceptibility-associated responses. In WT plants, NBR1-mediated selective autophagy targets the ABA-responsive transcription factor ABI5 and promotes its autophagic degradation, thereby restricting ABA signaling and limiting bacterial proliferation. In nbr1 mutants, impaired selective autophagy reduces ABI5 turnover, leading to ABI5 accumulation, enhanced expression of ABA-responsive genes, and increased susceptibility to bacterial infection. Thus, NBR1-mediated turnover of ABI5 acts as a regulatory mechanism that buffers ABA-associated susceptibility responses under warming conditions.

## ACKNOWLEGMENTS

We are grateful to Professor Jigang Li (College of Biological Sciences, China Agricultural University) for providing the pSuper:ABI5–3×Flag and pSuper:ABI5–GFP constructs. We thank Dr. Germán Robert (UDEA, CONICET-INTA, Argentina), Dr. Georgina Fabro (CIQUIBIC, CONICET-UNC, Argentina), and Dr. Elina Welchen (Instituto de Agrobiotecnología del Litoral, CONICET–UNL, Argentina) for providing plant materials used in this study.

## AUTHOR CONTRIBUTIONS

LAR, JS, and ILL designed the research and performed most of the experiments. AMYS contributed to bacterial growth assays and nucleic acid extraction. LAR and VLL generated and genotyped transgenic and mutant lines. MGT and VSF performed phytohormone quantification by LC-MS/MS. LAR and ILL analyzed the data. HRL and ILL conceived and supervised the study. ILL, NMC and HRL wrote the manuscript with contributions of LAR, AMYS and VLL. All authors revised the manuscript, approved the final version, and agree to be accountable for the work.

## SUPPLEMENTARY FIGURES

**Supplemental Figure 1. Autophagic flux is induced in 7-day-old Arabidopsis seedlings upon *Pma* infection and warming.** Seven-day-old 35S:GFP–ATG8a seedlings were flooded with mock solution (10 mM MgCl₂) or *Pma* at 1 × 10□ CFU ml⁻¹ and subsequently incubated at 22 °C or 29 °C for 1 day. **(a)** Autophagic flux was assessed by immunoblot detection of GFP and GFP–ATG8a and quantified by densitometric analysis as the ratio of free GFP to GFP–ATG8a. Each biological replicate consisted of five pooled seedlings per treatment. **(b)** Representative confocal images showing GFP–ATG8a-labeled autophagic structures and the corresponding quantification of autophagic puncta per image field in root and shoot tissues. For each treatment, three independent biological replicates were analyzed, and three images were acquired per replicate. Each circle represents an individual plant. Boxes indicate the median and interquartile range, and whiskers represent the minimum and maximum values. Arrowheads indicate representative GFP–ATG8a puncta used for quantification.

**Supplemental Figure 2. Warming enhances infection-induced autophagy independently of type III effector delivery.** 35S:GFGP-ATG8a plants were infiltrated with mock solution, *Pseudomonas syringae* pv. *tomato* DC3000 (*Pst*), or the type III secretion-deficient mutant *Pst-*Δ*hrcC*, at 5 × 10□ CFU ml⁻¹, and subsequently incubated at 22 °C or 29 °C. Autophagic flux was assessed at 2 days post-inoculation by immunoblot detection of free GFP and GFP–ATG8a, and band intensities quantification by densitometric analysis of the free GFP/GFP–ATG8a ratio. A representative immunoblot and the corresponding quantification are shown. Each circle represents an individual plant. Boxes indicate the median and interquartile range, and whiskers represent the minimum and maximum values. Asterisks indicate statistically significant differences compared to mock (*P < 0.05; two-way ANOVA followed by false discovery rate correction using the Benjamini, Krieger and Yekutieli two-stage linear step-up procedure).

**Supplemental Figure 2. Warming enhances infection-induced autophagy independently of type III effector delivery.** 35S:GFGP-ATG8a plants were infiltrated with mock solution, *Pseudomonas syringae* pv. *tomato* DC3000 (*Pst*), or the type III secretion-deficient mutant *Pst-*Δ*hrcC*, at 5 × 10□ CFU ml⁻¹, and subsequently incubated at 22 °C or 29 °C. Autophagic flux was assessed at 2 days post-inoculation (dpi) by immunoblot detection of free GFP and GFP–ATG8a, and quantified as the free GFP/GFP–ATG8a ratio. A representative immunoblot and the corresponding densitometric quantification are shown. Each circle represents an individual plant. Boxes indicate the median and interquartile range, and whiskers represent the minimum and maximum values. Asterisks indicate statistically significant differences compared to mock (*P < 0.05; two-way ANOVA followed by false discovery rate correction using the Benjamini, Krieger and Yekutieli two-stage linear step-up procedure).

**Supplemental Figure 3. NBR1 is required for tolerance to *Pseudomonas cannabina pv. alisalensis* under warming.** WT and *nbr1* plants were infiltrated with *Pma* at 5 × 10□ CFU ml⁻¹, and subsequently incubated at 22 °C or 29 °C. Bacterial growth was quantified at 2 days post-inoculation. **(a)** Bacterial titers expressed as CFU per cm² of leaf tissue. **(b)** Susceptibility index (SI), calculated as the ratio between bacterial titers at 29 °C and the mean bacterial titer at 22 °C for the same genotype. **(c)** Chlorophyll a and b concentrations. Each circle represents the mean value obtained from an independent experiment. In (a) and (b), boxes indicate the median and interquartile range, and whiskers represent the minimum and maximum values. In (c), bars represent the mean ± SE. Asterisks indicate statistically significant differences (*P < 0.05, **P < 0.01, ***P < 0.001) based on linear mixed-effects models including genotype and temperature as fixed factors and experiment as a random factor, followed by planned pairwise comparisons of estimated marginal means.

**Supplemental Figure 4. NBR1-dependent tolerance to bacterial infection operates in basal immunity but is dispensable during AvrRpt2-mediated ETI.** WT and *nbr1* plants were infiltrated with *Pseudomonas syringae* pv. *tomato* DC3000 (*Pst*), the type III secretion–deficient mutant *Pst-*Δ*hrcC*, or *P. syringae* pv. *tomato* expressing AvrRpt2 (*Pst-AvrRpt2*), at 5 × 10□ CFU ml⁻¹, and subsequently incubated at 22 °C or 29 °C. Bacterial growth was quantified at 2 days post-inoculation. **(a)** Bacterial titers expressed as CFU per cm² of leaf tissue at 22 °C and 29 °C. **(b)** Susceptibility index (SI), calculated as the ratio between bacterial titers at 29 °C and the mean bacterial titer at 22 °C for the same genotype and bacterial strain. Each circle represents an individual plant. Boxes indicate the median and interquartile range, and whiskers represent the minimum and maximum values. Asterisks indicate statistically significant differences (\**P* < 0.05, \*\*\**P* < 0.001; Kruskal–Wallis test for (a) and two-way ANOVA for (b), followed by false discovery rate correction using the Benjamini, Krieger and Yekutieli two-stage linear step-up procedure).

**Supplemental Figure 5.** Expression of salicylic acid- and abscisic acid-associated genes in WT and *nbr1* plants during bacterial infection under warming conditions. WT and *nbr1* plants were infiltrated with mock solution or *Pma* at 5 × 10□ CFU ml⁻¹, and subsequently incubated at 22 °C or 29 °C. Relative transcript levels of the SA-associated genes (a) *PR1* and (b) *ICS1*, and the ABA-associated genes (c) *NCED3*, (d) *ABI5*, (e) *SGR1* and (f) *NYC1*, normalized to *PP2A-A3*, and measured at 1 dpi. Each circle represents an individual plant. Boxes indicate the median and interquartile range, and whiskers represent the minimum and maximum values. Asterisks indicate statistically significant differences (*P < 0.05; two-way ANOVA or Kruskal–Wallis test followed by false discovery rate correction using the Benjamini, Krieger and Yekutieli two-stage linear step-up procedure).

**Supplemental Figure 6. ABI5–Flag co-immunoprecipitates with NBR1 in Arabidopsis seedlings.** Fourteen-day-old seedlings expressing pSuper:ABI5–Flag in WT and *atg5* backgrounds, or BIM1–Flag (Liang et al. 2018) as a negative control, were grown at 22 °C. Protein extracts were immunoprecipitated using anti-NBR1 magnetic beads. (a) Input and (b) immunoprecipitated fractions are shown. Red arrows indicate ABI5–Flag and BIM1–Flag proteins.

**Supplemental Figure 7. Basal turnover of ABI5–GFP in mock-treated plants.** pSuper:ABI5–GFP plants in WT, *nbr1* and *atg5* backgrounds were incubated at 22 °C or 29 °C for 1 day. ABI5 turnover was assessed by immunoblot detection of GFP and ABI5–GFP using an anti-GFP antibody and quantified by densitometric analysis of the free GFP/ABI5–GFP ratio. Representative immunoblots and the corresponding quantification are shown. Each circle represents an individual plant. Boxes indicate the median and interquartile range, and whiskers represent the minimum and maximum values. Statistical significance was evaluated using ANOVA on log₂-transformed ratios followed by false discovery rate correction using the Benjamini, Krieger and Yekutieli two-stage linear step-up procedure. Asterisks indicate statistically significant differences (***P < 0.001) relative to WT.

**Supplemental Figure 8. PCR genotyping of *nbr1 abi5* double mutants.** Genomic DNA from WT, *nbr1 abi5*, and *nbr1 abi5* plants was analyzed by PCR using gene-specific (RP+LP) and T-DNA border primers (LB) to confirm the presence of the corresponding mutant alleles. Representative amplification products are shown.

