## Supplemental Figure for "Selective autophagy promotes bacterial immunity under warming through NBR1-dependent regulation of ABI5"

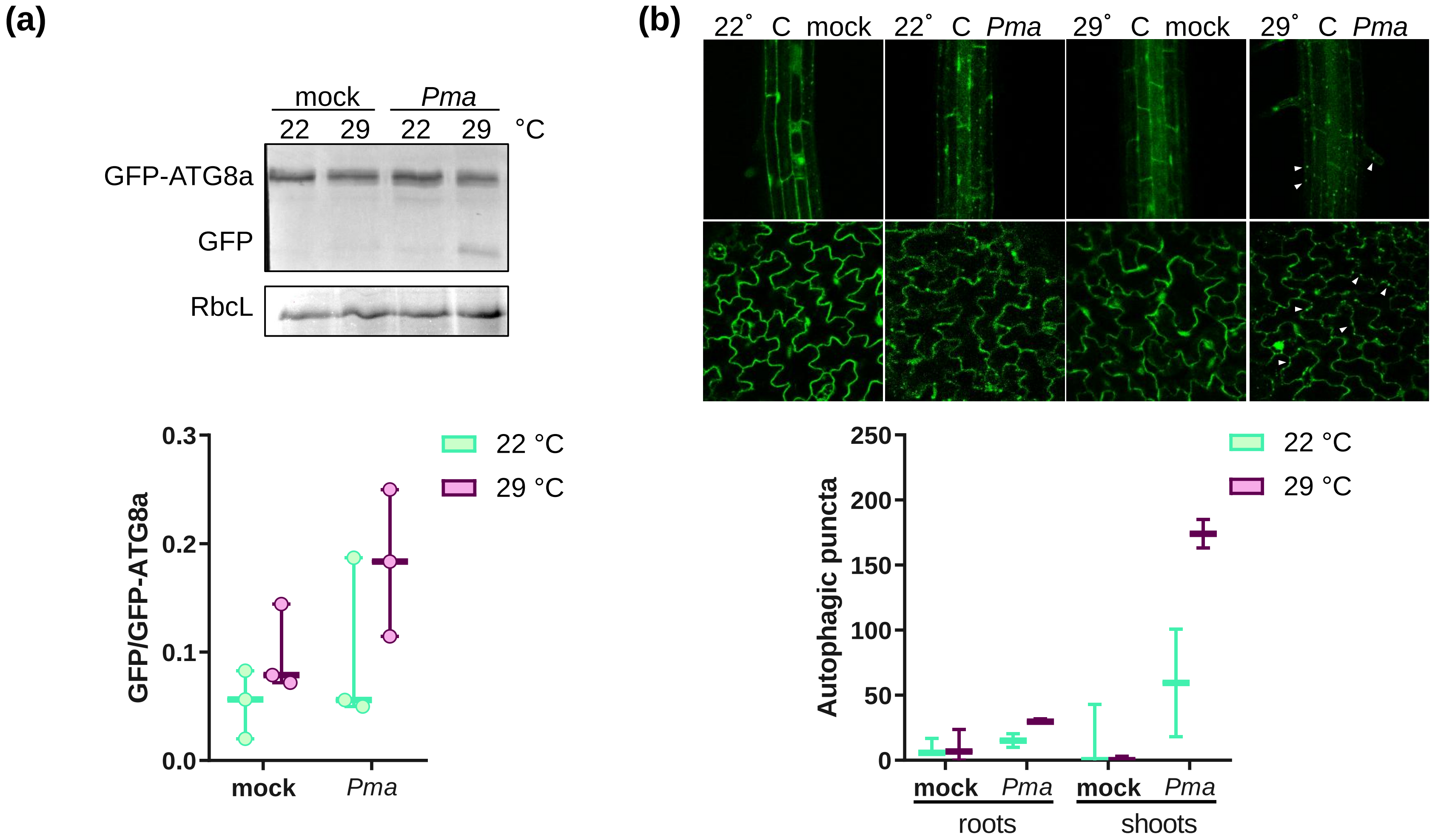


**Supplemental Figure 1. Autophagic flux is induced in 7-day-old Arabidopsis seedlings upon *Pma* infection and warming.** Seven-day-old 35S:GFP–ATG8a seedlings were flooded with mock solution (10 mM $MgCl_2$) or *Pma* at 1 × $10^7$ CFU $ml^{-1}$ and subsequently incubated at 22 °C or 29 °C for 1 day. **(a)** Autophagic flux was assessed by immunoblot detection of GFP and GFP–ATG8a and quantified by densitometric analysis as the ratio of free GFP to GFP–ATG8a. Each biological replicate consisted of five pooled seedlings per treatment. **(b)** Representative confocal images showing GFP–ATG8a-labeled autophagic structures and the corresponding quantification of autophagic puncta per image field in root and shoot tissues. For each treatment, three independent biological replicates were analyzed, and three images were acquired per replicate. Each circle represents an individual plant. Boxes indicate the median and interquartile range, and whiskers represent the minimum and maximum values. Arrowheads indicate representative GFP–ATG8a puncta used for quantification.

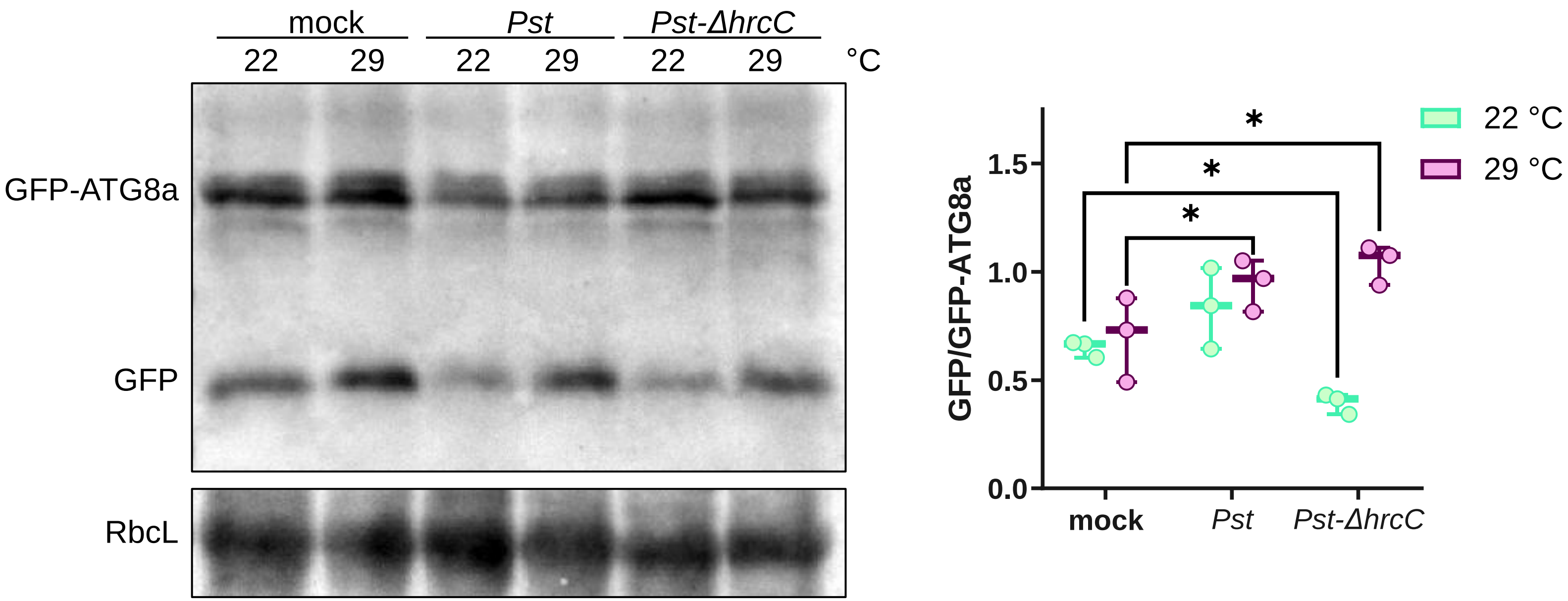


**Supplemental Figure 2. Warming enhances infection-induced autophagy independently of type III effector delivery.** 35S:GFGP-ATG8a plants were infiltrated with mock solution, *Pseudomonas syringae* pv. *tomato* DC3000 (*Pst*), or the type III secretion-deficient mutant Pst-ΔhrcC, at 5 × $10^5$ CFU $ml^{-1}$, and subsequently incubated at 22 °C or 29 °C. Autophagic flux was assessed at 2 days post-inoculation by immunoblot detection of free GFP and GFP–ATG8a, and band intensities quantification by densitometric analysis of the free GFP/GFP–ATG8a ratio. A representative immunoblot and the corresponding densitometric quantification are shown. Each circle represents an individual plant. Boxes indicate the median and interquartile range, and whiskers represent the minimum and maximum values. Asterisks indicate statistically significant differences compared to mock (*$P < 0.05$; two-way ANOVA followed by false discovery rate correction using the Benjamini, Krieger and Yekutieli two-stage linear step-up procedure).

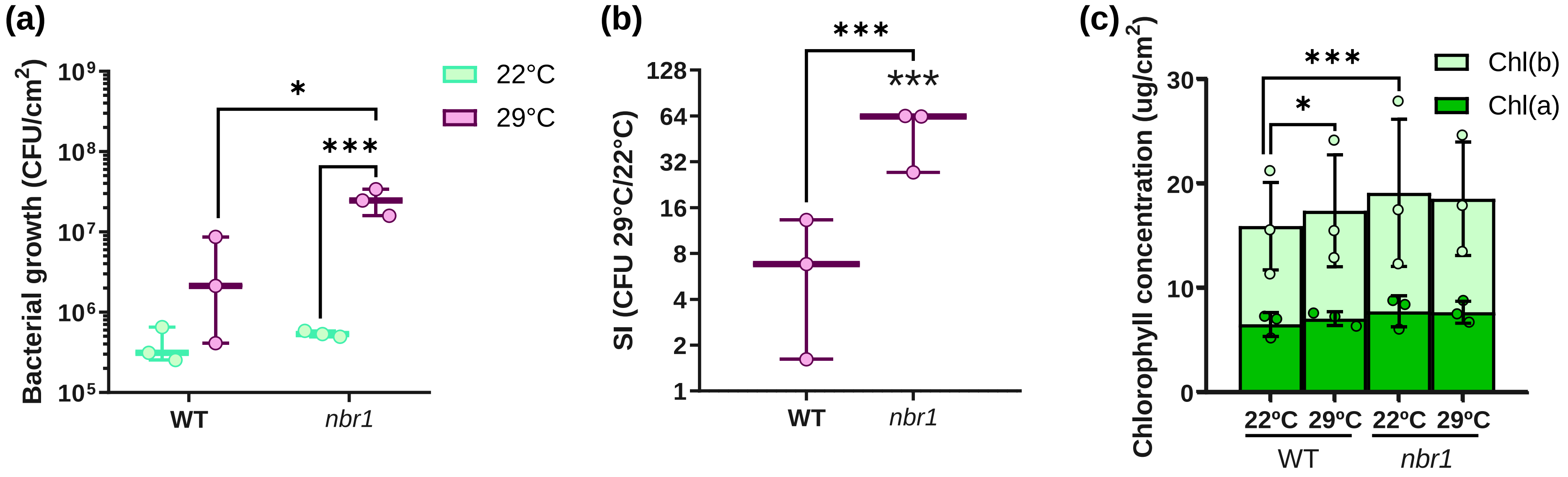


**Supplemental Figure 3. NBR1 is required for tolerance to *Pseudomonas cannabina pv. alisalensis* under warming.** WT and *nbr1* plants were infiltrated with *Pma* at 5 × $10^5$ CFU $ml^{-1}$, and subsequently incubated at 22 °C or 29 °C. Bacterial growth was quantified at 2 days post-inoculation. **(a)** Bacterial titers expressed as CFU per $cm^2$ of leaf tissue. **(b)** Susceptibility index (SI), calculated as the ratio between bacterial titers at 29 °C and the mean bacterial titer at 22 °C for the same genotype. **(c)** Chlorophyll a and b concentrations. Each circle represents the mean value obtained from an independent experiment. In (a) and (b), boxes indicate the median and interquartile range, and whiskers represent the minimum and maximum values. In (c), bars represent the mean ± SE. Asterisks indicate statistically significant differences (*P < 0.05, **P < 0.01, ***P < 0.001) based on linear mixed-effects models including genotype and temperature as fixed factors and experiment as a random factor, followed by planned pairwise comparisons of estimated marginal means.

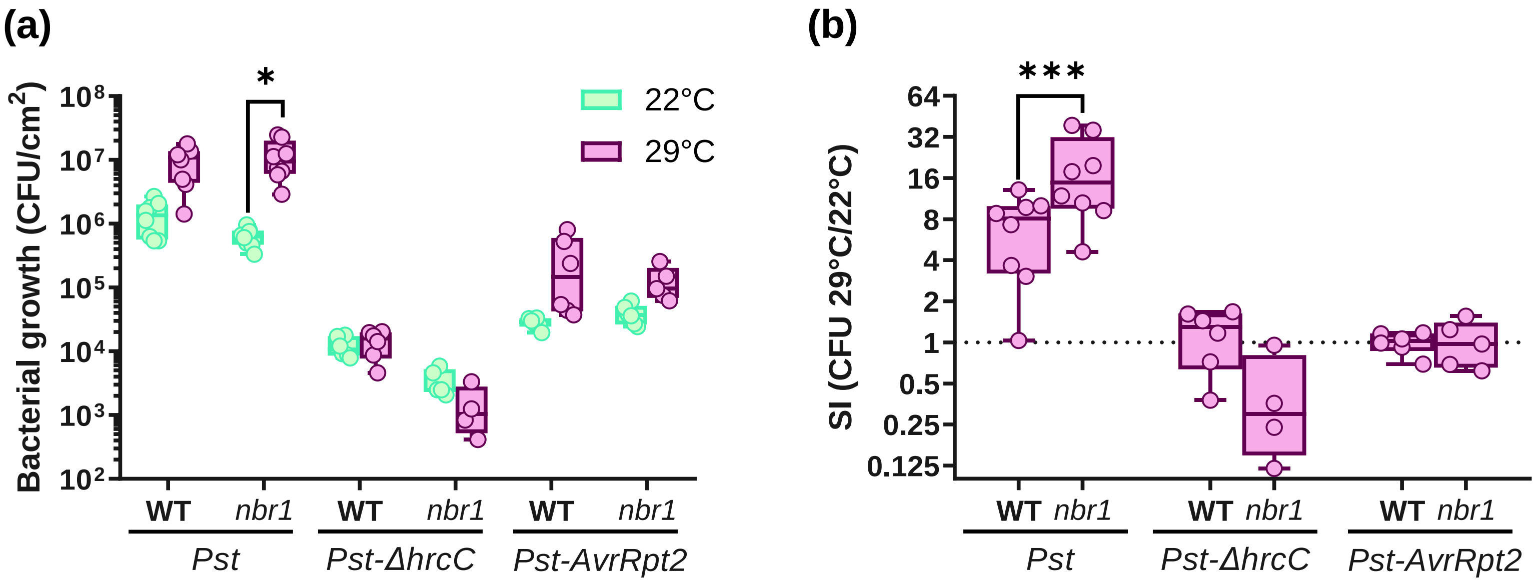


**Supplemental Figure 4. NBR1-dependent tolerance to bacterial infection operates in basal immunity but is dispensable during AvrRpt2-mediated ETI.** WT and *nbr1* plants were infiltrated with *Pseudomonas syringae* pv. *tomato* DC3000 (*Pst*), the type III secretion–deficient mutant Pst-ΔhrcC, or *P. syringae* pv. *tomato* expressing AvrRpt2 (*Pst-AvrRpt2*), at $5 \times 10^5$ CFU $ml^{-1}$, and subsequently incubated at 22 °C or 29 °C. Bacterial growth was quantified at 2 days post-inoculation. **(a)** Bacterial titers expressed as CFU per $cm^2$ of leaf tissue at 22 °C and 29 °C. **(b)** Susceptibility index (SI), calculated as the ratio between bacterial titers at 29 °C and the mean bacterial titer at 22 °C for the same genotype and bacterial strain. Each circle represents an individual plant. Boxes indicate the median and interquartile range, and whiskers represent the minimum and maximum values. Asterisks indicate statistically significant differences (*$P < 0.05$, ***$P < 0.001$; Kruskal–Wallis test for (a) and two-way ANOVA for (b), followed by false discovery rate correction using the Benjamini, Krieger and Yekutieli two-stage linear step-up procedure).

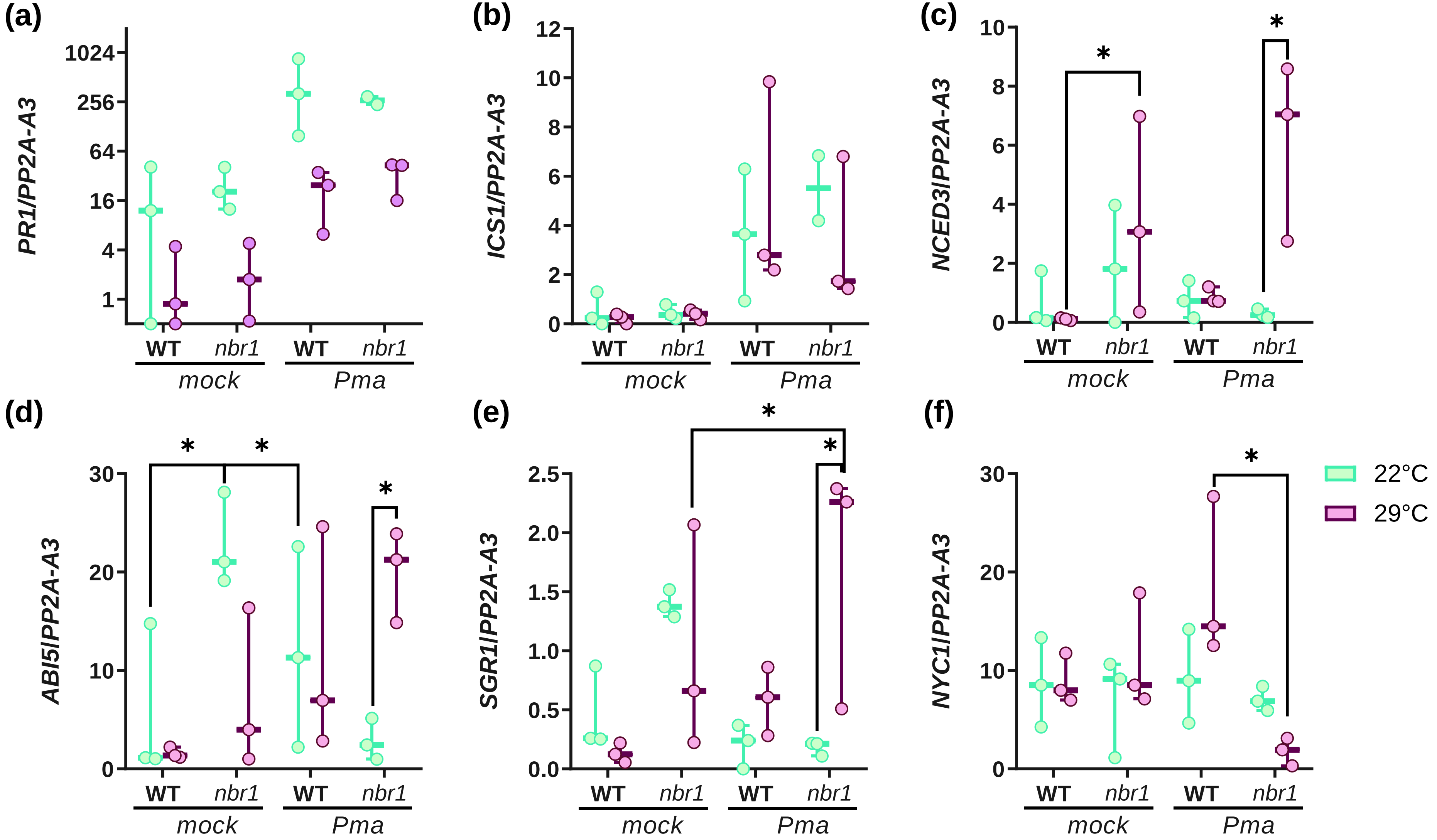


**Supplemental Figure 5. Expression of salicylic acid- and abscisic acid-associated genes in WT and *nbr1* plants during bacterial infection under warming conditions.** WT and *nbr1* plants were infiltrated with mock solution or *Pma* at 5 × $10^5$ CFU $ml^{-1}$, and subsequently incubated at 22 °C or 29 °C. Relative transcript levels of the SA-associated genes **(a)** *PR1* and **(b)** *ICS1*, and the ABA-associated genes **(c)** *NCED3*, **(d)** *ABI5*, **(e)** *SGR1* and **(f)** *NYC1*, normalized to *PP2A-A3*, and measured at 1 dpi. Each circle represents an individual plant. Boxes indicate the median and interquartile range, and whiskers represent the minimum and maximum values. Asterisks indicate statistically significant differences (**P* < 0.05; two-way ANOVA or Kruskal–Wallis test followed by false discovery rate correction using the Benjamini, Krieger and Yekutieli two-stage linear step-up procedure).

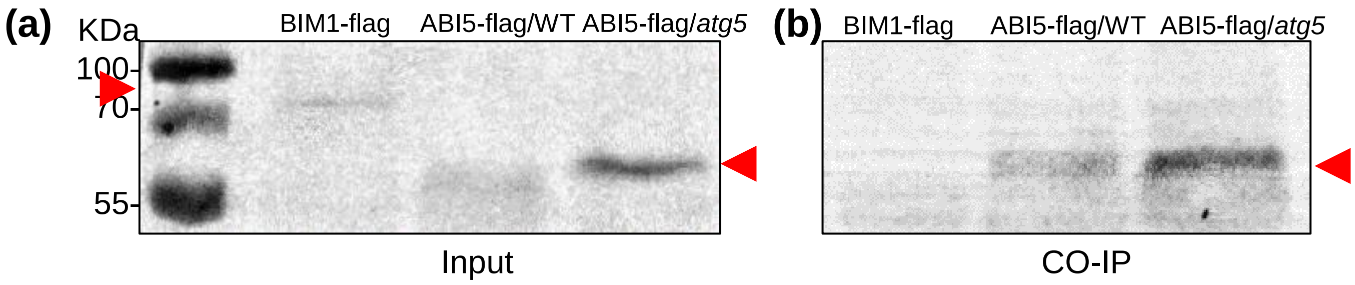


**Supplemental Figure 6. ABI5–Flag co-immunoprecipitates with NBR1 in Arabidopsis seedlings.** Fourteen-day-old seedlings expressing pSuper:ABI5–Flag in WT and *atg5* backgrounds, or BIM1–Flag (Liang et al. 2018) as a negative control, were grown at 22 °C. Protein extracts were immunoprecipitated using anti-NBR1 magnetic beads. (a) Input and (b) immunoprecipitated fractions are shown. Red arrows indicate ABI5–Flag and BIM1–Flag proteins.

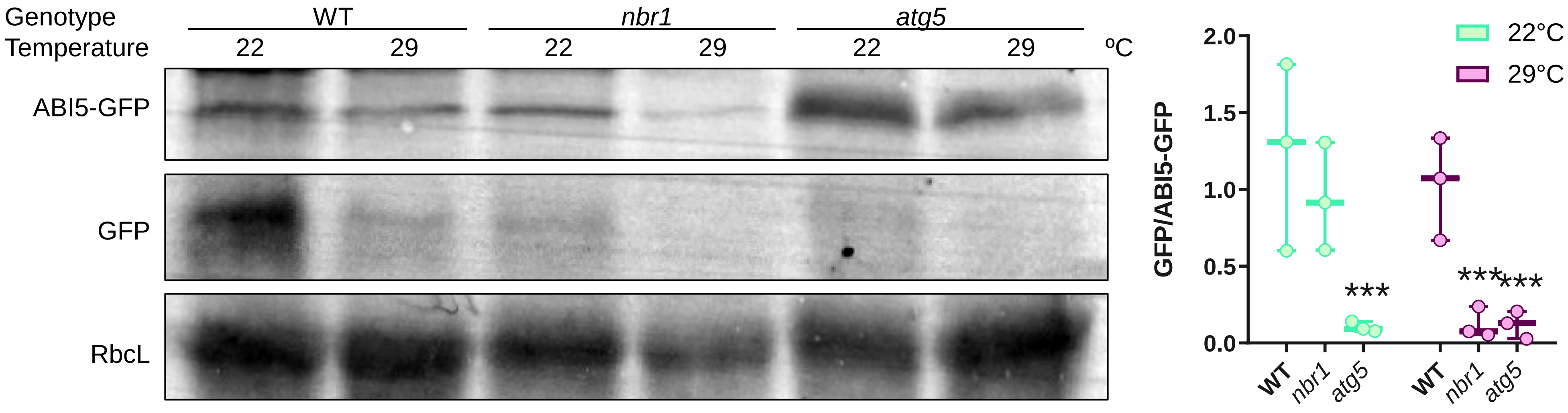


**Supplemental Figure 7. Basal turnover of ABI5–GFP in mock-treated plants.** pSuper:ABI5–GFP plants in WT, *nbr1* and *atg5* backgrounds were incubated at 22 °C or 29 °C for 1 day. ABI5 turnover was assessed by immunoblot detection of GFP and ABI5–GFP using an anti-GFP antibody and quantified by densitometric analysis of the free GFP/ABI5–GFP ratio. Representative immunoblots and the corresponding quantification are shown. Each circle represents an individual plant. Boxes indicate the median and interquartile range, and whiskers represent the minimum and maximum values. Statistical significance was evaluated using ANOVA on $\log_2$-transformed ratios followed by false discovery rate correction using the Benjamini, Krieger and Yekutieli two-stage linear step-up procedure. Asterisks indicate statistically significant differences (***P < 0.001) relative to WT.

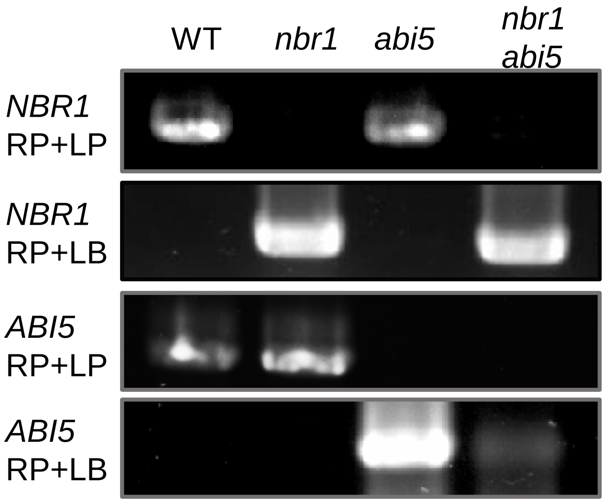


**Supplemental Figure 8. PCR genotyping of *nbr1 abi5* double mutants.** Genomic DNA from WT, *nbr1 abi5*, and *nbr1 abi5* plants was analyzed by PCR using gene-specific (RP+LP) and T-DNA border primers (LB) to confirm the presence of the corresponding mutant alleles. Representative amplification products are shown.
